# Single-nucleus multiple-organ chromatin accessibility mapping in the rat

**DOI:** 10.1101/2024.11.11.622900

**Authors:** Ronghai Li, Yue Yuan, Shanshan Duan, Qiuting Deng, Wen Ma, Chang Liu, Peng Gao, Li Lu, Chuanyu Liu

## Abstract

The chromatin accessibility landscape is the basis of cell-specific gene expression. We generated a multiorgan, single-nucleus chromatin accessibility landscape from the model organism *Rattus norvegicus*. For this single-cell atlas, we constructed 25 libraries via snATAC-seq from nine organs in the rat, with a total of over 110,000 cells. Cell classification integrating gene activity scores with known marker genes identified 77 cell types, which were strongly correlated with those in published mouse single-cell transcriptome atlases. We further investigated the enrichment of cell type- and organ-specific transcription factors (TFs), the dynamics of T-cell developmental trajectories across organs, and the conservation and specificity of gene expression patterns across species. These findings provide a foundation for further investigations of the cell composition and gene regulatory networks throughout the rat body.

**Highlights:**

- Generation of a single-cell atlas of chromatin accessibility in nine organs of the rat
- Characterization of cell type- and organ-specific transcription factors (TFs)
- Dynamics of chromatin accessibility in developing T cells revealed by cross-organ analysis
- Conservation and specificity of gene expression patterns among humans, mice, and rats revealed by cross-species analysis

## Introduction

The Human Cell Atlas (HCA) project aims to create a comprehensive reference cell atlas of all cells in the human body (the basic unit of life). This will serve as a basis for understanding human health and for diagnosing, monitoring and treating disease. However, a key scientific question is what insights can be gained from cell atlases. To date, studies of cell atlases have advanced our understanding of anatomy, development, physiology, pathology, and intra- and intercellular regulation at a new level of granularity. They have also advanced our understanding of cellular diversity, revealing the cellular compositions of complex tissues and organs and how cells interact with each other in states of health and disease^1^. The development of single-cell and spatial genomics technologies, as well as the corresponding algorithms, has enabled the mapping of cells across omics^2,3^, organs^4^, species^5^, developmental states^6^, and diseases^7^ with unprecedented resolution. This has facilitated the systematic probing of biological questions related to cell type, spatial location^8^, developmental trajectory^9^, fate determination^10^, the tumour microenvironment^11^, and molecular mechanisms, among others. These advances provide powerful new tools and will open new avenues for clinical medicine, especially in precision medicine and personalized treatment.

The rat (*Rattus norvegicus*) has long played a key role in scientific research as a model animal for studies of disease mechanisms, drug development and physiology. In recent years, the development and application of single-cell technology has facilitated considerable advances in the mapping of single cells within organs in the rat. For example, single-cell atlases of individual organs (e.g., the brain^12,13^, kidney^14^, and testes^15^) have provided valuable insights into the cellular compositions and gene regulatory mechanisms of these organs. Furthermore, such studies are progressively expanding to produce more systemic single-cell atlases encompassing multiple organs^16^, elucidating the cellular heterogeneity and synergistic interactions between different organs. In parallel, cross-species single-cell mapping analyses of individual organs are also being conducted. For example, recent studies utilizing scRNA-seq of lung tissues derived from mice, rats, pigs and humans have revealed evolutionarily conserved mechanisms in alveolar intercellular communication^17^. Nevertheless, a comprehensive multiorgan, single cell epigenomic profile of the rat has yet to be established, with the understanding of chromatin accessibility in the rat lagging considerably behind that in other species. The capture of intracellular chromatin-accessible regions is a fundamental aspect of determining the mechanisms through which genes are regulated in diverse cell types and states.

To address this disparity, we employed snATAC-seq to map chromatin accessibility in nine organs of the rat, thereby effectively addressing a previously existing research gap in this area and providing a valuable resource for the scientific community. We systematically analysed more than 110,000 nuclei captured from nine organs, identified 77 cell types, identified hundreds of thousands of candidate *cis*-regulatory elements (cCREs) and characterized cell type-specific and organ-specific transcription factors (TFs). Overall, these data reveal the epigenetic landscapes of major organs in the rat at an unprecedented resolution, thereby providing valuable information for understanding the functions and interactions of cells within organs. Furthermore, the creation of this resource enables cross-species comparative studies and enhances our comprehension of the similarities and distinctions between mammalian cell types and gene regulatory mechanisms.

## Results

### Single-nucleus multiple-organ chromatin accessibility mapping in the rat

Here, we report the cellular composition and chromatin accessibility landscape of multiple organs in the rat. The atlas consists of single-cell epigenomic data from 115,723 nuclei isolated from nine organs (namely, the thyroid, thymus, heart, lung, liver, spleen, kidney, pancreas and ovary) of a single female Sprague‒Dawley rat aged 7–8 months (Figure 1A). The organs were dissociated into single-nuclear suspensions in accordance with preestablished methods and then subjected to snATAC-seq via the standard MGI DNBelab C4 scATAC-seq protocol (STAR Methods). A total of 25 libraries were generated, with two or three technical replicates performed for each organ.

**Figure 1.**
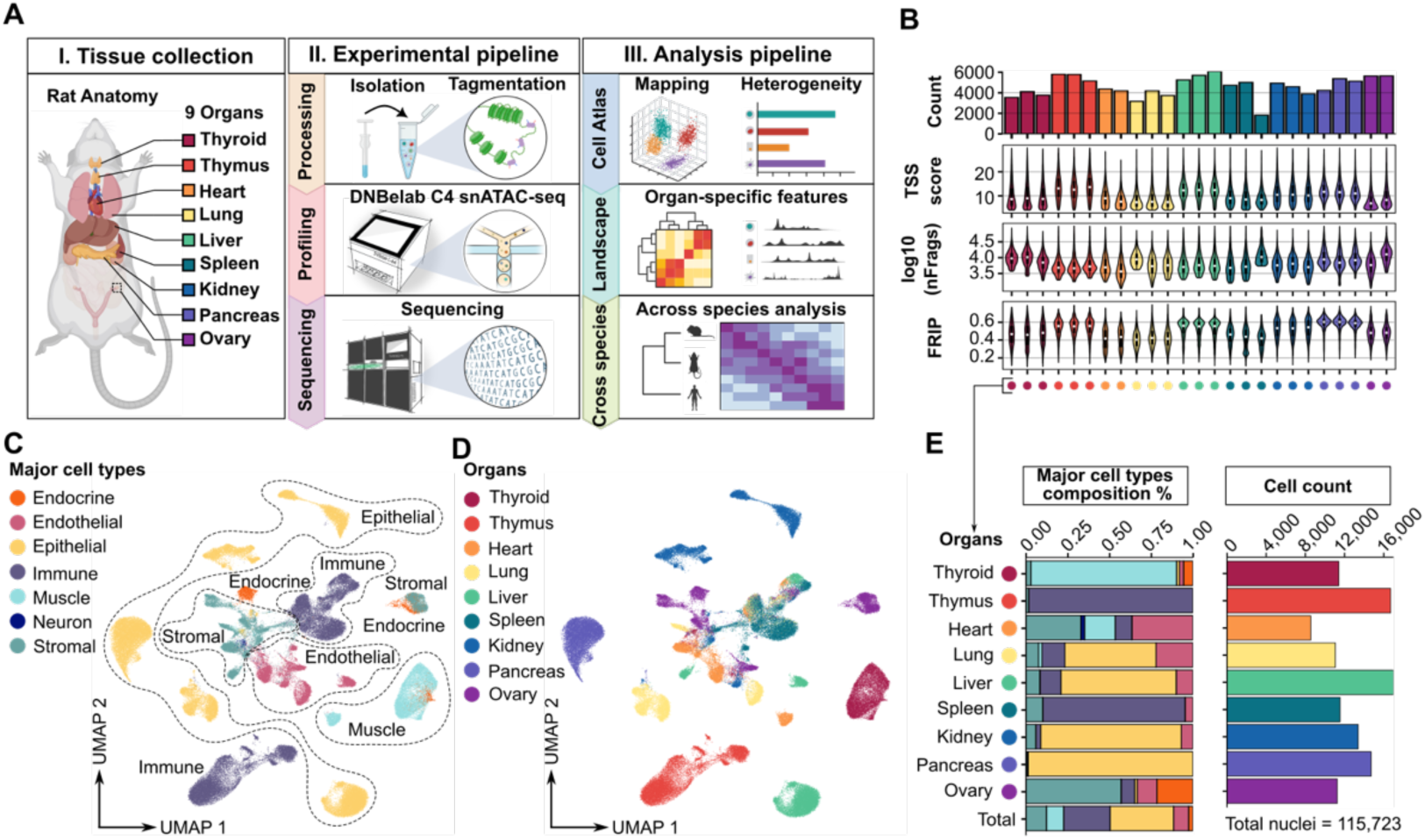
Cross-organ single-nucleus ATAC sequencing atlas of an adult rat. **A**. Schematic of the study design, including the tissue collection, experimental pipeline, and analysis pipeline steps (Created in BioRender.com). **B**. Quality control metrics across the library, with the same colour used to represent technical replicates of the same organ. The bar plot shows the number of nuclei in each library. Violin plots show the transcription start site (TSS) enrichment scores, log10 unique nuclear fragment counts, and fraction of reads in peaks (FRIP) for each library. **C**. Cross-organ snATAC-seq atlas. The UMAP plot shows all the nuclei identified in this study, coloured according to major cell type. **D**. Same as C but coloured according to organ. **E**. Stacked bar plot showing the fraction of major cell types in each organ, with the total proportions of major cell types in this study shown in the bottom column (left). The bar plot shows the number of nuclei for each organ, and the total number of nuclei in this study is shown at the bottom (right).

To obtain high-quality single-cell profiles, we applied a three-step data filtering operation to the raw data (STAR Methods). First, the initial filtering process excluded approximately 28,274 cells with a TSS less than 4 and unique nuclear fragments per cell less than 1,000, which are typically regarded as low-quality cells (Figures S1A and S1B). Next, 6,638 potential doublets, typically situated between clusters (Figures S1C and S1D), were filtered out. These doublets were predicted with an algorithmic approach that involved mixing the reads from thousands of individual cell combinations to synthesize simulated duplexes, which are most likely to be doublets. Then, the dataset was divided by organ of origin for data quality control and cell annotation. This approach was adopted to avoid batch effects between organs, thereby enhancing the accuracy of the annotation. Additionally, it enabled the identification of organ-specific cell types, thus improving the reliability of the annotation. In this phase, the low-quality cell clusters, comprising 3,571 cells, were defined in accordance with the established metrics and removed (Figure S1E). Ultimately, 115,723 high-quality cells were obtained (Figures 1B and S1F-S1H).

To define cell types, we analysed each organ independently by performing iterative LSI-based dimensionality reduction on the genome-wide tile matrix of all cells, followed by shared nearest neighbour (SNN) modularity optimization-based clustering (STAR Methods). To define clusters of the snATAC dataset accurately, we employed a methodology based on the derivation of gene scores through modelling of the assessment of chromatin accessibility near a specific gene in ArhcR, with the goal of inferring expression level of the gene. This process integrates signals from the entire gene body and scales the signal with bidirectional exponential decay from the gene TSS and the gene transcription termination site, while accounting for neighbouring gene boundaries^18,19^. We combined known cell type-specific expressed marker genes (Table S1) with genes differentially expressed between clusters in this dataset (Data S3) to comprehensively assess and assign their cell subtype labels (Figures S2A-S2I; STAR Methods). The major cell types of the clusters were manually categorized on the basis of the defined cell subtypes, and the labels were transferred back to the full dataset. The marker genes of the major cell types were visualized to assess the accuracy of the global clustering across organs and the relationships between cells from different organs (Figure S1I). Overall, we identified 7 major cell types: endothelial, endocrine, muscle, immune, endothelial, stromal and neuron (Figure 1C). Endothelial cells expressed *Cdh1*, *Krt18* and *Krt8*. *Star*, *Cyp19a1* and *Cyp11a* are expressed in the endocrine system. Muscles expressed *Acta1*, *Myh7* and *Myh1*. Immune cells expressed *Cd3d*, *Cd4* and *Cd163*. Endothelial cells expressed *Flt1*, *Pecam1* and *Vmf*. Stromal cells expressed *Dcn*, *Col1a1* and *Col3a1*. We identified neurons with high expression of *Scn7a*, *Grik3*, *Ntng1*, *Gfra2* and *Ncam1* only in the heart (Figure S1I). Notably, examination of the gene expression profiles of neurons in the hearts of mice and humans in publicly available datasets revealed that *Grik3*, *Ntng1*, and *Gfra2* were highly expressed exclusively in the rat heart dataset^20,21^. This unexpected finding suggests that our data provide a wealth of new information about the rat chromatin accessibility landscape.

To visualize differences in the chromatin accessibility landscape across organs, we employed UMAP to visualize all cells and differentiate their colours according to their respective cellular origins (Figure 1D; Data S1 and S2). Stromal cells, immune cells, and endothelial cells from different organs tend to be clustered together (i.e., by cell type) rather than clustered according to the organ of origin or sample batch (Figure 1E). This phenomenon has been identified in previously published data^22,23^ and may emphasize the commonality of certain cell types in different organs. Furthermore, in accordance with expectations, we observed that immune cells are the major cell types of the thymus and spleen, which are the primary immune organs in vivo. However, these cells tended to cluster by organ rather than by cell type (Figure 1E). This phenomenon was also observed in epithelial cells from multiple organs, suggesting that the chromatin accessibility of these cells is distinctly organ specific. For example, immune cells situated within the thymus are immature, whereas those located within the spleen are mature^24^. The epithelial cells of each organ display distinct morphological, gene expression and functional characteristics in accordance with their environmental and functional contexts^4^.

### Cross-organ cell type identification and comparison with mouse single-cell RNA sequencing data

In addition to utilizing gene scores for the purpose of assigning cluster identity, we can reference published scRNA datasets to facilitate the identification of cluster identity for the snATAC dataset (STAR Methods). A total of 8 publicly available mouse scRNA datasets were screened for matching organ origins (Figure 2A). However, none of the datasets matched thyroid samples, which again emphasizes the value of our dataset in providing novel biological information. We integrated the scRNA dataset with the snATAC dataset for each organ individually via cellular alignment (Figure 2B). This method employs unsupervised identification of pairs of cells with similar biological states (defined as anchors) between datasets, followed by the joint projection of the features of the two modalities into a shared low-dimensional space^25^. To provide a more intuitive assessment of the results of data integration, the predicted score was used to evaluate the accuracy and confidence of integration between cells, whereas the Jaccard index was employed to assess the correlation between transferred labels of the RNA cell subtype (automated annotation) and the labels of the ATAC cell subtype that were manually annotated on the basis of gene scores (manual annotation) (Figures S3A– S3H). In total, 77 cell subtypes were identified (Figure 2C), and a high degree of correspondence was observed between the automatic and manual cell type identification methods (Figure 2D). This not only validates and enhances the reliability of the dataset for cell type annotation but also allows further exploration of similarities and differences between mice and rats for cell type annotation in the same organ.

**Figure 2.**
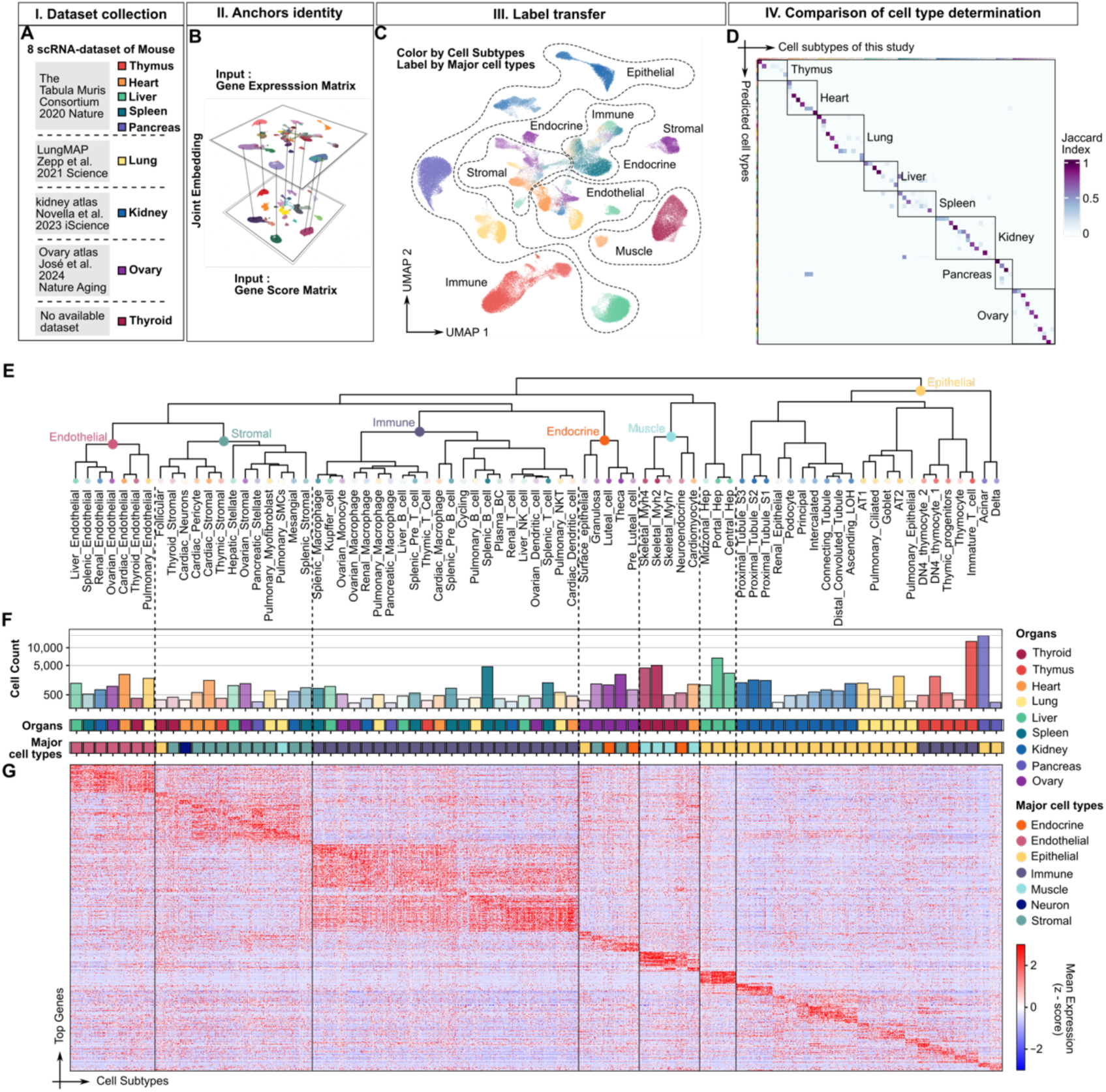
Cross-organ cell type identification and comparison via single-cell RNA sequencing data. **A**. The table lists the scRNA datasets for eight mouse organs, including the thymus, heart, liver, spleen, pancreas, lung, kidney, and ovary. No dataset is available for the thyroid. **B.** A 3D plot showing the jointly embedded gene expression matrix and gene score matrix to identify anchor points across the datasets. **C**. UMAP plot showing the locations of cells, with cells coloured by subtype and labelled by major cell type. **D**. The Jaccard index was used to quantify the overlap between automated annotation and manual annotation. **E**. Hierarchical clustering dendrogram illustrating the relationships and similarities between the cell subtypes across organs, with coloured labels corresponding to the major cell type categories. **F**. Bar plot showing the number of nuclei for each cell subtype. The cell subtype labels are aligned with the dendrogram in E. Below the cell count bar plot, the first colour bar represents the organs from which the cell subtypes were sourced, and distinct colours are used for each organ. The second colour bar indicates the classification of the cell subtypes into major cell types. The legend indicating the colour codes for both the organs and major cell types is located on the right side of the plot. **G.** Heatmap showing the expression patterns of the top genes across the different cell subtypes identified in the study. The x-axis corresponds to the cell subtypes, ordered as in dendrogram E, whereas the y-axis lists the top genes, with their expression levels colour-coded from low (blue) to high (red).

The major goal of creating a cross-organ cell atlas is to gain a comprehensive understanding of cell type diversity and their relationships between different organs. We employed the gene score matrix to cluster average gene expression profiles from 77 cell subtypes, with the objective of exploring similarities and specificities in chromatin accessibility across cells in different organs (Figure 2E).

The same major cell types from different organs tend to cluster together, such as endothelial cells, stromal cells, epithelial cells, and immune cells (Figure 2F). This phenomenon is also observed in more finely categorized subpopulations, such as the clustering of macrophages, B cells, and T cells among the immune cells. This finding indicates a comparable pattern of chromatin accessibility and gene activity for these cell types across different organs, as demonstrated by our heatmap of gene expression profiles mapped in accordance with the cell type ordering of the hierarchical clustering results (Figure 2G). These findings suggest that these genes are subject to some degree of conservation in terms of their functional characteristics and regulatory mechanisms. Furthermore, despite originating from different organs, the trillions of cells in the body originate from a single cell, namely, the fertilized egg. Thus, it would be reasonable to hypothesize that cell types associated with a similar developmental sequence would cluster together with a higher degree of similarity. This is an interesting question worthy of further investigation. Furthermore, the application of hierarchical clustering serves to validate the reliability of cellular annotations, indicating that the similarity and specificity among cell types have been successfully captured. This is based on the premise that cell types of the same broad class from different organs will tend to cluster together.

Despite the observation that cells of the same types from disparate organs tend to cluster, we noted that certain cell types exhibit organ-specific clusters, a phenomenon that is particularly evident in epithelial cells, such as those of the liver, kidney, and lung (Figure 2F). Although these epithelial cells share certain fundamental features, such as *Cdh1*, *Krt18*, and *Krt8* expression (Figure S1I), they exhibit notable organ specificity. Epithelial cells in the liver (e.g., hepatocytes) specifically express *Arg1*, *Gls2*, and *Cyp2e1* (Figure S2E), exemplifying their functions in amino acid metabolism, energy metabolism, and detoxification^26^. Epithelial cells in the kidney (e.g., proximal tubules) specifically express *Slc3a1*, *Cubn*, *Slc34a1*, *Slc7a13*, and *Slc5a1* (Figure S2G), reflecting their unique functions in substance transport, nutrient absorption, and maintenance of homeostasis in the body^27^. Epithelial cells in the lung (e.g., alveolar type 1/2 cells) specifically express *Emp2* and *Lrrk2* (Figure S2D), consistent with their multiple functions in maintaining lung homeostasis, performing gas exchange, repairing damage and regulating immune responses^28^. These organ-specific functional requirements produce unique chromatin accessibility patterns in epithelial cells in different organs, which is reflected in the organ-based clustering observed in hierarchical clustering analyses.

In summary, our findings emphasize two fundamental cellular relationships between different organs: cell type specificity (similarity across organs) and organ specificity (similarity within the same organ). The same major cell types (e.g., immune cells, endothelial cells and stromal cells) display comparable chromatin accessibility profiles across different organs, indicating that these cells share common functional attributes in diverse tissues. In contrast, organ specificity denotes the environmental adaptation and functional differentiation of cells within a specific organ (e.g., epithelial cells), resulting in a heightened degree of similarity between different cell types within the same organ.

### Characterization of specific TF motifs across cell types and organs in the rat

TFs play crucial regulatory roles in organ development, cell type differentiation and maintenance of function. We aggregated chromatin accessibility data from the same major cell types into ’pseudobulk replicates’ to improve the signal‒to‒noise ratio for peak calling (STAR Methods), thereby facilitating more accurate identification of approximately 450,000 open chromatin regions that represent potential *cis*-elements, primarily promoters, introns, exons, and distal regulatory regions (such as enhancers) (Figure 3A). To identify cell subtype-specific TFs, we employed motif enrichment analysis of chromatin open regions (peaks) specific to each cell subtype. This approach enabled us to ascertain which TF has regulatory functions within these regions. The heatmaps of the motif enrichment results for each cell subtype were subsequently sorted according to the previously described hierarchical clustering results (Figure 2E). The motif enrichment patterns were distributed predominantly according to cell type, indicating that TF activity can be employed to reconstruct known relationships and functional profiles between different cell types (Figures 3B and 3C; Data S4). For example, *Sfpi1*^29,30^, a TF linked to the formation of blood cells, is specifically enriched in immune cells (especially Kupffer cells, ovarian monocytes, and splenic macrophages), indicating that it plays a pivotal role in immune cell differentiation and function. *Esrrb*, a TF associated with stem cell development, pluripotency, and germline development, was specifically enriched in certain endocrine cells of the ovary (e.g., luteal, granulosa, and theca cells), indicating that it may be involved in the regulation of ovarian function. *Foxa1*^31^ and *Snai2*^32^, key regulators of epithelial cell differentiation, were specifically enriched in epithelial cell clusters (e.g., acinar, AT2, and central hepatocytes), reflecting their importance in maintaining epithelial cell characteristics. *Etv2*^33^, which is associated with angiogenesis and endothelial cell differentiation, was enriched not only in endothelial cells (e.g., liver and ovarian endothelial cells) but also in certain immune cells (e.g., splenic T cells and Kupffer cells), indicating that it may be involved in regulating the interaction between immune cells and the vascular system. These findings demonstrate that distinct TFs influence the differentiation and functional sustenance of cell types, thereby substantiating the intimate correlation between TF activity and cell fate and functional status.

**Figure 3.**
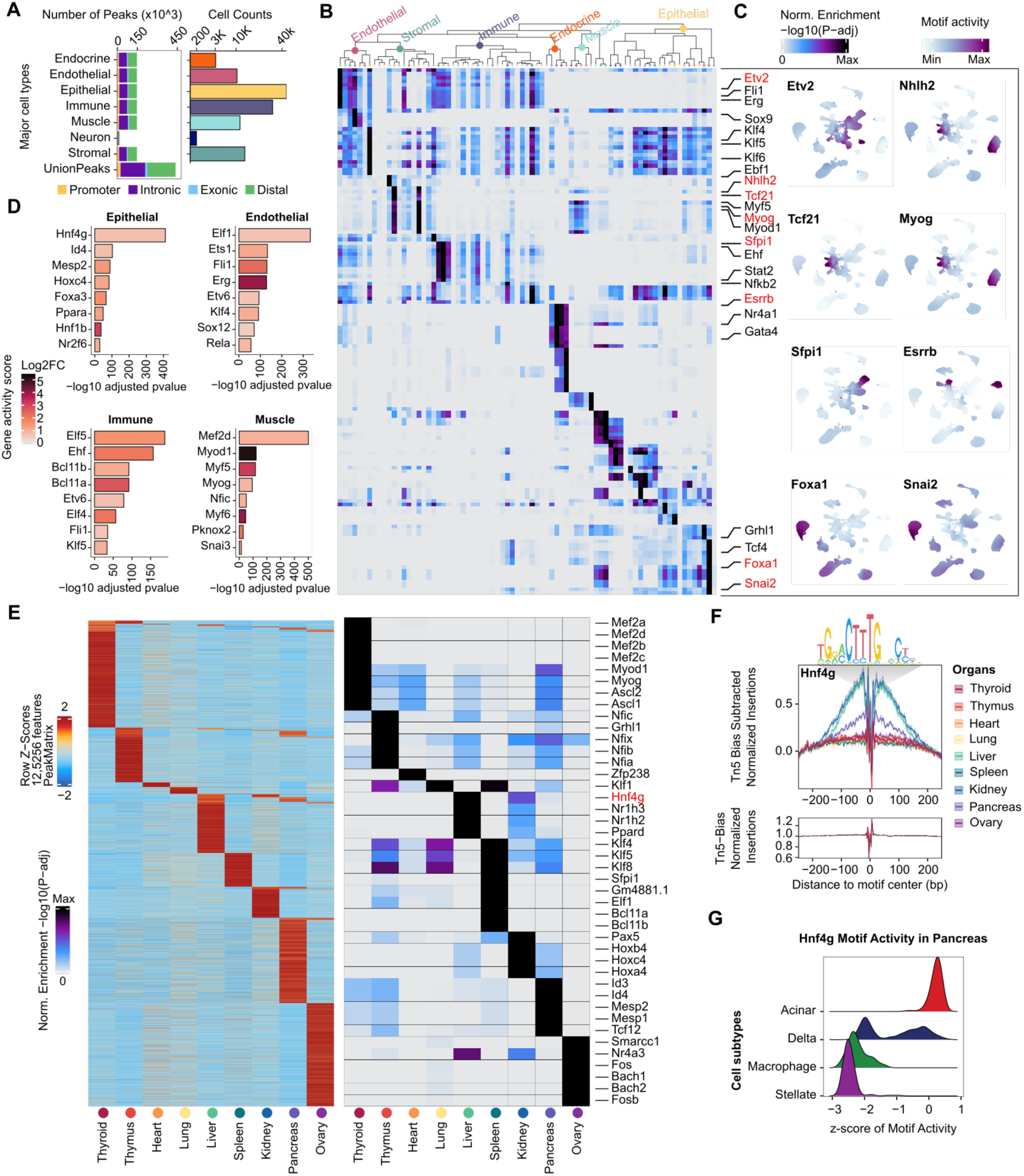
Characterization of specific TF motifs across cell types and organs in the rat. **A.** Bar plot (left) showing the number of chromatin accessibility peaks detected for different genomic features across major cell types. The bar plot (right) shows the number of nuclei for each major cell type. **B.** Hypergeometric enrichment of TF motifs in marker peaks for each cell subtype. The rows represent different TFs, and the TFs of interest are shown on the right. The columns represent the different cell subtypes and are arranged according to the hierarchical clustering dendrogram at the top. **C**. UMAP plots showing TF motif activity across the cellular landscape. **D**. Bar plots showing the top 8 TFs (significance of differential motif activity scores) of each major cell type. The colour of the bars indicates the log2-fold change (log2FC) in the gene activity score, with darker red representing a higher degree of differential activity. **E**. The left heatmap displays the normalized enrichment (Z scores) of accessible chromatin regions (peaks) in different organs, and the right heatmap shows the enrichment of TF motifs in peaks in different organs, represented by -log10 adjusted p values (P-adj). Each row relates to a specific TF. **F.** TF footprint plot showing the enrichment of the TF *Hnf4g* across different organs. The top plot shows the sequence logo of the *Hnf4g* motif. The middle plot displays the Tn5 bias-subtracted normalized insertions centred on the *Hnf4g* motif, with different lines representing different organs and higher peaks representing greater chromatin accessibility. The bottom plot shows the Tn5 bias around the motif centre, confirming that the observed enrichment is not due to bias in Tn5 transposase activity. **G.** Ridge plot showing the distribution of *Hnf4g* motif activity (measured as z scores) across different pancreatic cell types.

To identify and visualize TFs associated with chromatin accessibility in different major cell types, we combined the gene scores with motif enrichment data to elucidate the link between the activity of specific TFs and gene regulatory potential (Figure 3D; Data S5; STAR Methods). Although this approach does not reflect gene expression levels directly, it reveals the relationship between chromatin accessibility and potential regulators, thereby providing important information regarding the specific TFs in different cell types.

To determine which TFs may be primarily responsible for establishing and maintaining organ-specific regulatory programs, we employed motif enrichment analysis of peaks that were specific to each organ (STAR Methods).

We characterized numerous TFs as either shared or specific to individual organs (Figure 3E; Data S6). One example is *Hnf4g*, which is a class of nuclear receptor TFs and member of the *HNF4* family. It has been shown to be expressed predominantly in organs that are involved in metabolic processes, such as the liver^34^, kidney^35^, and intestine^36^. Notably, slight enrichment of *Hnf4g* was observed in the pancreas (Figure 3E). To confirm the presence of specific binding sites in the pancreas, TF footprinting was employed, which revealed that although the binding strength was weaker than that in the liver and kidney, *Hnf4g* indeed has specific binding sites in the pancreas (Figure 3F). Recently, *Hnf4g* was identified as an oncogene in pancreatic cancer, and it has been demonstrated to regulate cell proliferation and migration in pancreatic cancer^37,38^. Nevertheless, it remains unclear which pancreatic cell types are involved in this process. Consequently, we investigated the enrichment of *Hnf4g* in distinct pancreatic cell types and discovered high basal activity of *Hnf4g* in pancreatic acinar cells (Figure 3G), indicating that *Hnf4g* may play a pivotal regulatory role in the physiological processes of these cells. These findings also indicate that future studies should focus on the regulatory network and signalling pathways of *Hnf4g* in acinar cells to gain a deeper understanding of its role in pancreatic cancer.

In summary, our dataset offers a comprehensive map of the TF motifs in different organs, major cell types and cell subtypes in the rat. This information will facilitate the elucidation of gene regulatory networks in various organs and cell types and the identification of the potential roles of TFs in cell function and fate determination and provide a crucial foundation for subsequent basic research and disease studies.

### Diversity of immune cells across organs and accessible chromatin changes associated with T-cell lineage commitment

As previously stated, the construction of a single-cell atlas across organs in rats revealed the presence of diverse cell types and varying degrees of chromatin accessibility in different organs. This atlas was constructed to help us understand the spatial heterogeneity observed among organs. From an ontological perspective, all the organs of an intact individual are derived from a single cell. Therefore, over time, cells execute a series of processes, including division, development, differentiation, and maturation, which ultimately lead to the formation of distinct cell types and states or apoptosis. This implies that even within a single temporal frame, there is heterogeneity in cells at different stages of development. This motivated us to attempt to resolve the single-cell atlas from a temporal perspective, with the aim of elucidating how cells gradually acquire specific functions and what regions of chromatin are activated or repressed during differentiation. Here, we investigated the interrelationships among immune cells across organs, with a particular emphasis on the developmental trajectory of T cells.

We repeated dimensionality reduction, batch correction, and clustering with the cells defined as immune cells from the full dataset and performed peak calling and motif enrichment analyses at a finer level within groups of cell subtypes (STAR Methods). The selection of an appropriate clustering resolution is a pivotal aspect of single-cell data analysis. Different resolutions result in the identification of varying numbers and granularities of cell populations, which directly impacts the characterization and biological interpretation of cell subpopulations. In this study, we utilized Clustree^39^ to examine and contrast the clustering outcomes at varying resolutions to gain insight into the dataset structure and identify suitable clustering parameters. Clustree was initially developed for the analysis of single-cell RNA data. However, in this study, we applied it to single-cell ATAC data, as there was no significant difference between the two in terms of data structure (STAR Methods). A total of 14 clustering resolutions were selected for investigating the population relationships of immune cell populations at varying resolutions. If we disregard the biological implications and consider population delineation exclusively in terms of the data structure, it appears that the number of cell subpopulations that can be identified increases concomitantly with increased clustering resolution (Figure S4A).

Importantly, however, as the clustering resolution increases, more clusters with smaller proportions and new clusters consisting of multiple parent clusters begin to appear (Figure S4B). These are often artificially segmented by algorithmic transition clusters that do not have distinct biological significance or unique gene expression/chromatin accessibility features. In accordance with these principles, a clustering resolution of 0.5 was selected as it is stable in terms of clustering relationships and does not produce a considerable number of fringe groups for the division of the immune cell population (Figure 4A). Surprisingly, the results of the population delineation presented essentially a one-to-one correspondence with the cell subpopulations that had been originally defined by organ clustering alone (Figure 4B). This finding not only validates that clustering at this resolution effectively captures the differences between cell populations but also illustrates the accuracy of our cell type definitions.

**Figure 4.**
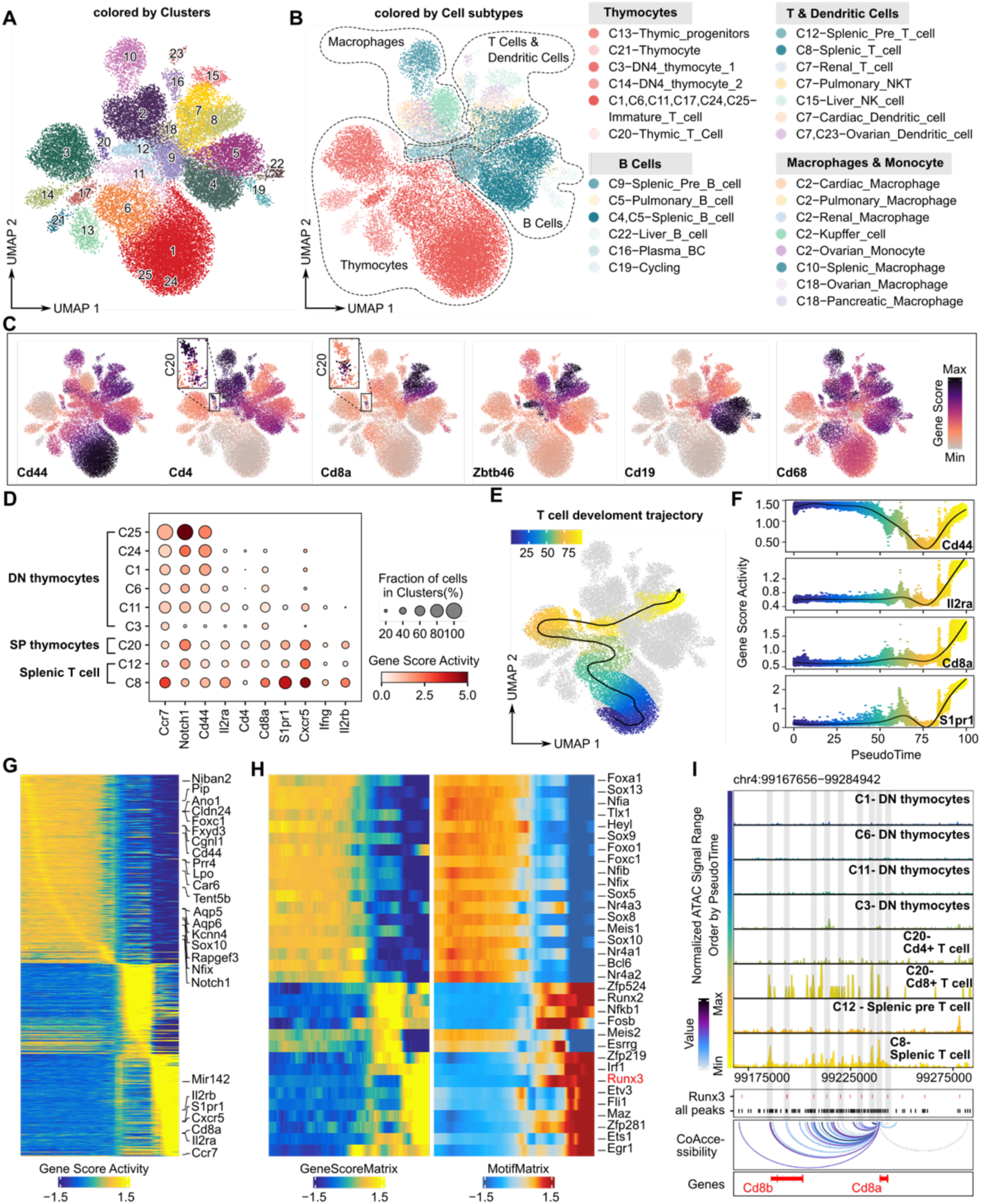
Diversity of immune cells across organs and accessible chromatin changes associated with T-cell lineage commitment. **A.** UMAP plot showing the clustering of single cells from various organs, coloured according to distinct clusters (C1 to C25). **B.** UMAP plot showing the cell subtypes of immune cells in different organs labelled by major cell type. **C**. UMAP plot showing the gene activity scores of known marker genes in major cell types. **D**. Dot plot showing the gene activity score levels of known marker genes across distinct clusters annotated as T cells. The dot size corresponds to the percentage of cells expressing the genes in each cluster, and the colour represents the average expression level. **E**. UMAP plot showing the T-cell development trajectory, with cells coloured according to pseudotime. **F.** The plots display gene score activities for key genes during pseudotime, highlighting dynamic changes in chromatin accessibility as T cells differentiate. **G.** Heatmap showing the gene activity score changes along the T-cell development pseudotime trajectory; the top genes are shown on the right. **H.** Paired heatmaps of positive TF regulators whose TF motif activity (right) and matched gene activity score (left) are positively correlated across the T-cell development pseudotime trajectory. **I**. Genomic tracing of *Cd8a* accessibility over pseudotime during T cell development. Selected TFs are highlighted in the feature tracks using different colours. Loops below the tracks show coaccessibility interactions between genomic regions, highlighting potential regulatory interactions that may drive T-cell development.

We classified immune cells into four main groups: thymocytes; B cells; T cells and dendritic cells; and macrophages and monocytes, which each originate from multiple organs (e.g., the kidneys, liver, heart, ovaries and spleen) (Figure 4B). The different groups can be classified on the basis of the gene scores of known marker genes. For example, dendritic cells specifically express *Zbtb46*, B cells specifically express *Cd19*, cytotoxic T cells specifically express *Cd8a*, and macrophages and monocytes express *Cd68* (Figure 4C). Notably, the C20 cluster, which has been defined as thymic T cells, appears to comprise two distinct cell subpopulations. One subpopulation presented high expression of *Cd4*, whereas the other presented high expression of *Cd8a* (Figure 4C). This observation may be indicative of the process of T-cell differentiation from the double-positive to the single-positive stage within the thymus. This motivated us to attempt to resolve the chromatin accessibility alterations that occur during T-cell development across different organs. The initial step was to delineate T cells at different developmental stages via gene expression profiles of gene score proxies (Figure 4D). Briefly, haematopoietic progenitors enter the thymus in the presence of the chemokine receptor *Ccr7* and then initiate T-cell lineage formation in response to Notch signalling. *Cd44* and *Il2ra* are expressed at the DN stage, *Cd4* and *Cd8a* are expressed at the DP and SP stages, respectively, and the expression of *Ccr7* and *S1pr1* is upregulated in preparation for exit from the thymus. The expression of *Ccr7* and *S1pr1* is maintained in primary T cells, which facilitates T-cell homing in the lymph nodes and spleen. The expression of *Cd44*, *Ifng*, *Il2rb* and *Cxcr5* is increased in activated and memory T cells, which have adapted to their functional requirements^40^. Notably, the definition of T cells at the DP stage was not included in this study, primarily because the thymus samples were collected from adult individuals, who are expected to have a lower number of thymic DP cells than a developing foetus or a young individual would^41^. To identify the regulatory elements that drive T-cell development, we subsequently reconstructed T-cell developmental trajectories and identified numerous fluctuations in chromatin accessibility, which significantly overlapped with existing findings (Figure 4E‒4G). To identify positive TF regulators, we combined the gene score matrix with the motif matrix for a comprehensive analysis (STAR Methods). In total, 33 positive TF regulators were identified along the T-cell developmental trajectory (Figure 4H). It was observed that the chromatin accessibility of *Runx3* became increasingly accessible at a later stage of the trajectory. *Runx3*, a pivotal factor in determining the fate of CD8+ T cells during T-cell differentiation, promotes *Cd8a* and *Cd8b* expression to ensure that CD4+ T-cell differentiation signals are inhibited^42^. To confirm the regulatory relationship of *Runx3* with the *Cd8a* gene, we visualized chromatin accessibility near the *Cd8a* gene, screened for coaccessible regions that overlap with only the promoter region of *Cd8a*, and highlighted potential individual binding sites for *Runx3* (STAR Methods). It was evident that the chromatin of the *Cd8a* promoter region was more accessible in Cd8+ T cells and splenic T cells and that this accessibility correlated strongly with that of the *Cd8b* promoter region (Figure 4I). Furthermore, numerous peaks co-opening with the *Cd8a* promoter region were annotated as *Runx3* binding sites. These findings suggest that these genomic regions are involved in the regulation of *Cd8a* expression as potential DNA binding sites of the TF *Runx3*. These observations lend further support to the hypothesis that *Runx3* may act as a positive regulator of the *Cd8a* gene.

In summary, we examined a cross-organ single-cell atlas from a temporal perspective to reconstruct T-cell developmental trajectories. The findings revealed many potential TFs that may regulate T-cell differentiation. This research has also provided insights into how to investigate the regulatory relationships between specific TFs and genes in greater depth.

### Cross-species analysis revealed similarities and differences in gene expression patterns in human, mouse and rat heart tissue

One crucial application for single-cell atlases is in cross-species analyses to gain insights into the origin and evolution of different organs and cell types, the conservation and specificity of species’ gene expression patterns^43,44^, and the identification of species-specific cell types^45^. However, many current cross-species integration algorithms were originally designed for use with scRNA datasets^46^. Furthermore, no standard integration method for scATAC datasets has been established in the field^47^. To further expand the applications of rat single-cell chromatin accessibility mapping, we attempted cross-omics and cross-species integration analyses to explore the conservation and species specificity of gene expression patterns in different organs (Figure 5A). The objective of this investigation was to ascertain whether gene scores could serve as proxies for molecular features in the context of cross-species dataset integration. To test this hypothesis, a dual-omics dataset comprising heart and kidney samples was analysed. In brief, highly variable homozygous genes were identified across species datasets and used as anchors for data integration via the Seurat V4 CCA method (STAR Methods). In the heart, we observed that cells from different datasets were effectively integrated, with high consistency in the clustering of the same cell types and in the gene expression levels of marker genes in the same cell type between species (Figure S5A-S5E). For example, CMs present a high degree of similarity in their marker genes across species. Similar results were observed in the kidney (Figure S5F-S5I). The results of our tests further emphasized that cell types are highly conserved across species in terms of certain important molecular mechanisms and functions. Additionally, we demonstrated that the use of gene scores as molecular features is a reliable strategy for cross-species integration of different omics data, thus further supporting the feasibility of comparative cross-species analyses.

**Figure 5.**
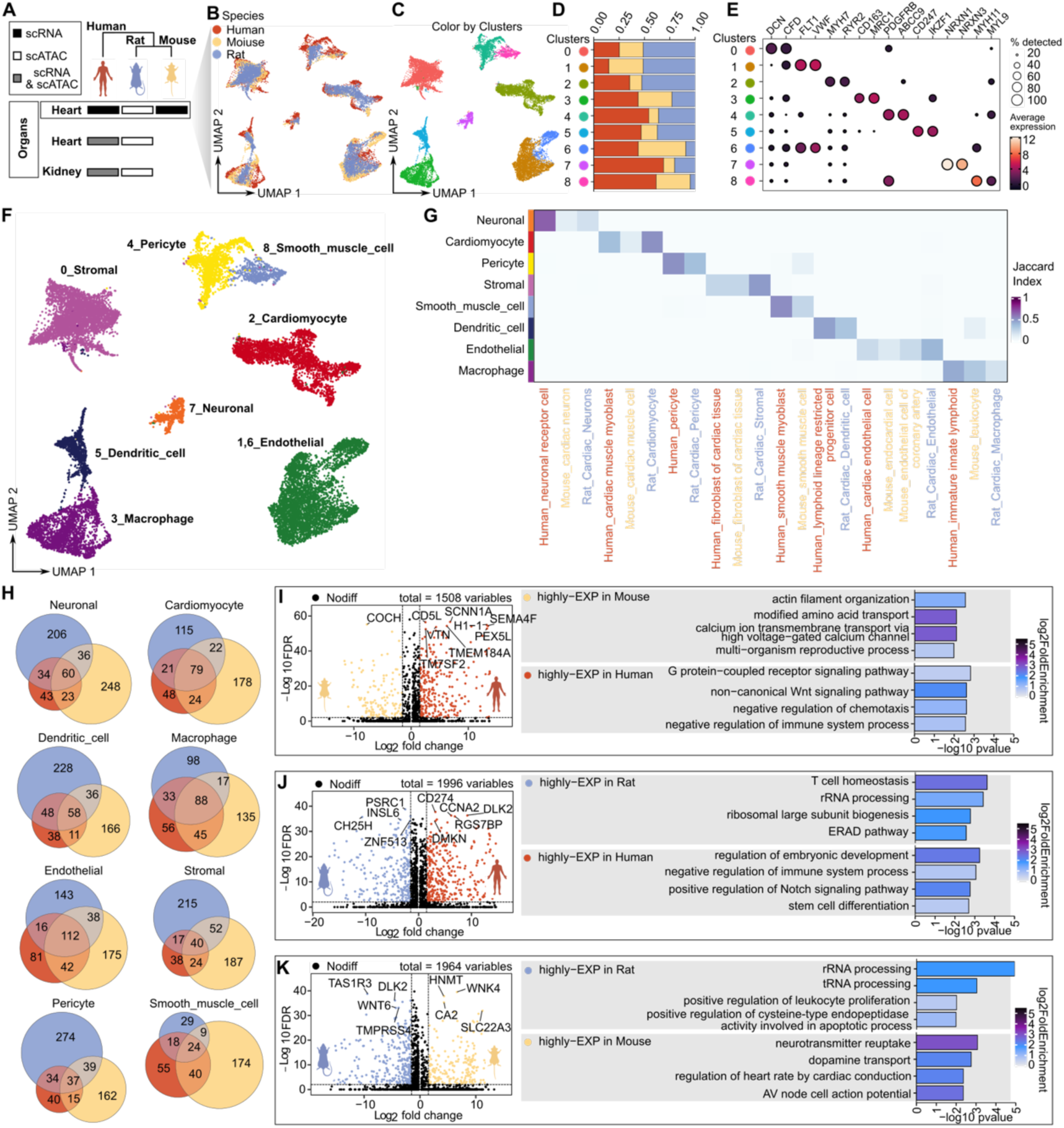
Comparative single-cell transcriptomic and epigenomic analysis across species. **A.** Overview of the types of data (scRNA-seq and scATAC-seq) collected from heart and kidney tissues across humans, rats, and mice. **B.** UMAP plot showing species-specific clustering. **C.** UMAP plot showing the clustering of single cells in the integrated dataset, coloured according to cluster. **D**. Stacked bar plot showing the fraction of each cluster across species (human, mouse, and rat). The different colours represent the different species, with the same colour scheme as in B. **E**. Dot plot displaying the gene expression levels of known marker genes of heart tissue across distinct clusters. The size of the dot corresponds to the percentage of cells expressing the genes in each cluster, and the colour represents the average expression level. **F.** UMAP plot showing the cell types of the integration dataset, coloured by cell type, from the manual annotation of the data in E.**G.** Heatmap showing the similarity (Jaccard index) of cell types across different species. The rows represent the manually annotated cell type labels in the cross-species integration dataset, and the columns represent the cell type label with which the cell was annotated in the original dataset. **H.** Venn diagrams showing the overlap of differentially expressed genes between species for specific cell types. **I**. Volcano plot showing the results of differential gene expression analysis between human and mouse cardiomyocytes. On the right side, the bar plot shows the top biological processes that are significantly enriched among the DEGs between species. **J**. Same as I but for the results of DEG analysis between human and rat cardiomyocytes. **K**. Same as I but for the results of DEG analysis between mouse and rat cardiomyocytes.

This integration strategy was subsequently applied to the integration of human, mouse, and rat datasets, and it was observed that cells between species exhibited a high degree of intermixing in each cluster (Figure 5B-5D). We manually annotated the major cell types in the integrated cardiac dataset on the basis of known gene markers and investigated discrepancies in cell type annotations between species (Figure 5E-5F). Our findings indicated that the major cell types were identified across species, which highlights the conservation of cell types between species (Figure 5G). Notably, however, cells annotated as pericytes in the human and rat datasets were annotated as smooth muscle cells in the mouse dataset. Similarly, cells annotated as lymphoid lineage-restricted progenitor cells in the human dataset were annotated as dendritic cells in the rat dataset and as leukocytes in the mouse dataset. We believe that this coclustering of cells assigned different markers in different datasets occurs largely due to differences in annotation granularity, as well as to differences in the dataset or analytical methodology on which the annotation is based. For example, the current human cardiac cell atlas includes 21 immune cell subpopulations^21^. To further explore the extent to which gene expression patterns are shared and differ across homologous cell types in different species, we performed cross-species identification of DEGs after downsampling the integrated dataset to 200 cells per cell type (STAR Methods). We observed a high degree of conservation of gene expression within the same cell type across species. For example, CMs presented 79 overlapping genes, macrophages presented 88 overlapping genes, and ECs presented 112 overlapping genes (Figure 5H; Data S7). However, most genes were enriched for expression in only one species, reflecting species specificity.

To further examine the species specificity of gene expression patterns, we concentrated our attention on cardiomyocytes, which exhibit relatively high conservation across species (STAR Methods). In general, genes that are more highly expressed in mice (such as *COCH*) are enriched in biological processes such as actin filament organization, amino acid transport, and calcium ion transmembrane transport. On the other hand, genes that are more highly expressed in humans (such as *CD5L*, *SCNN1A*, and *CD274*) are associated with pathways such as G protein-coupled receptor signalling, Wnt signalling, and immune system regulation (Figure 5I). Similarly, for the rat dataset, genes such as *PSRC1* and *DLK2* are associated with biological processes including T-cell homeostasis and leukocyte proliferation (Figure 5J and 5K). These cross-species comparisons underscore specific regulatory pathways that are more active in one species than in the other, illustrating the complexity of gene expression regulation across evolutionary contexts.

In summary, our results demonstrate that gene scores can be utilized as reliable proxies for gene expression in cross-species data integration analyses, with the capacity to discern biologically salient features across species. We employed this method to investigate the degree of conservation of cell types and the extent of shared and distinct gene expression profiles in human, mouse, and rat hearts. The results provide a valuable foundation for elucidating cross-species differences and similarities in heart biology while also offering insights into species-specific gene regulatory mechanisms.

## Discussion

The central dogma of molecular biology is that gene expression commences with the transcription of a DNA sequence, and chromatin accessibility has been identified as an important prerequisite for the regulation of gene expression. In recent years, single-cell analysis of transposase-accessible chromatin sequencing (scATAC-seq) has emerged as a powerful tool for elucidating regulatory patterns and cellular heterogeneity. Numerous exciting single-cell chromatin accessibility profiles have been generated via this tool, providing a robust foundation for the analysis of gene regulatory programs across tissues, developmental stages, and cell types within species^44,48–50^. However, the absence of a universal single-cell chromatin accessibility landscape for the rat (*Rattus norvegicus*), an experimental organism frequently employed in psychological, pharmacological, and behavioural studies, motivated us to generate a single-cell chromatin accessibility landscape across organs.

In this study, we used MGI DNBelab C4 scATAC-seq to create a single-cell atlas of chromatin accessibility from nine organs in the rat. This atlas of chromatin accessibility offers new insights into cellular regulation, uncovering gene activity in specific cell types and allowing systematic and comparative studies of cell types across various organs. The dataset comprises over 110,000 cells captured from nine organs, with 77 identified cell types and hundreds of thousands of cCREs. Additionally, we characterize cell type-specific and organ-specific TFs, offering detailed insights into the regulatory landscapes of various cell types and organs. This provides a valuable framework for exploring both normal biological processes and disease states.

Despite the similarity in chromatin accessibility observed between endothelial and stromal cells when examining the heterogeneity of cell types across different organs, we continued to assign distinct cell labels according to the organ of origin. This aspect is often overlooked in single-cell atlases from a single organ and, in our view, impedes cross-atlas comparisons. As single-cell technology has developed and cross-organ single-cell atlases have become more widely used, we anticipate that a consensus will be reached within the field to create a more standardized and uniform cell nomenclature.

The approach adopted in this study, which entailed dividing the organ dataset for cell annotation and subsequently integrating cell labels from disparate organs, proved more conducive to precise cell type definition. This was evidenced by the observation that immune cells from different organs tended to exhibit similar characteristics and cluster together while retaining some organ specificity. This has the effect of blurring the boundaries between cell types and increasing the difficulty of cell type definition. Furthermore, this method avoids the issue of batch effects among samples from different organs.

In the process of creating a single-cell atlas, the accurate definition of cell type and the cell state presents a significant challenge^51,52^ and is typically reflected in the choice of clustering resolution. With respect to this issue, we tested several clustering resolutions when exploring immune population heterogeneity in the present study. As the resolution increases, the number of cell populations that can theoretically be divided simultaneously also increases. Notably, there have been numerous discussions within the scientific community regarding the appropriate definition of biologically significant cell subclusters. We posit that the current approach, designated hicat^53^, represents a reasonable methodology for this purpose. This approach combines consensus clustering with iterative refinement through multiple rounds of high-variance gene selection, downscaling, dimensionality filtering, and clustering until no further subclusters satisfy the criteria for differential gene expression or cluster size termination. However, this method has certain limitations in terms of identifying rare populations and weak biological differences. The issue of defining cell types and cell states can be effectively addressed through the creation of comprehensive single-cell atlases that encompass the entire life cycle of an organism.

There are several limitations of this work that need to be considered. Notably, the data presented in this study were derived from a single rat. While this may limit the ability to capture rare cell types, our objective is to create a universal single-cell atlas from healthy rats. The identification of cell types is mainly dependent on sequencing depth, technology platforms, and data processing and analysis methods^54^. To increase the technical stability and reproducibility of the experimental data, multiple technical replicates were conducted for each organ. Second, as there is no comprehensive motif database dedicated to *Rattus norvegicus*, we employed *Mus musculus* (source CisBP) for peak annotations in this study. Given the high degree of genomic similarity between rats and mice, it can be assumed that the two should be relatively consistent. Third, in cross-species studies, our analyses concentrated on homologous genes and shared cell types, which may have resulted in the omission of some species-specific gene regulatory and expression patterns. Furthermore, we have not yet been able to identify cell types that are unique to a single species.

In summary, we constructed a comprehensive single-nucleus multiple-organ chromatin accessibility map in the rat, which will serve as a valuable resource for investigating gene regulation and functional studies across organs, species, and disease models. Furthermore, numerous aspects of our dataset have yet to be fully elucidated. These include comparisons of other cell lineages (e.g., endothelial, epithelial and stromal) across organs, as well as multiomics integration analyses (e.g., combining single-cell transcriptomes, DNA methylation, histone modifications, and other multiomics data).

## STAR★Methods

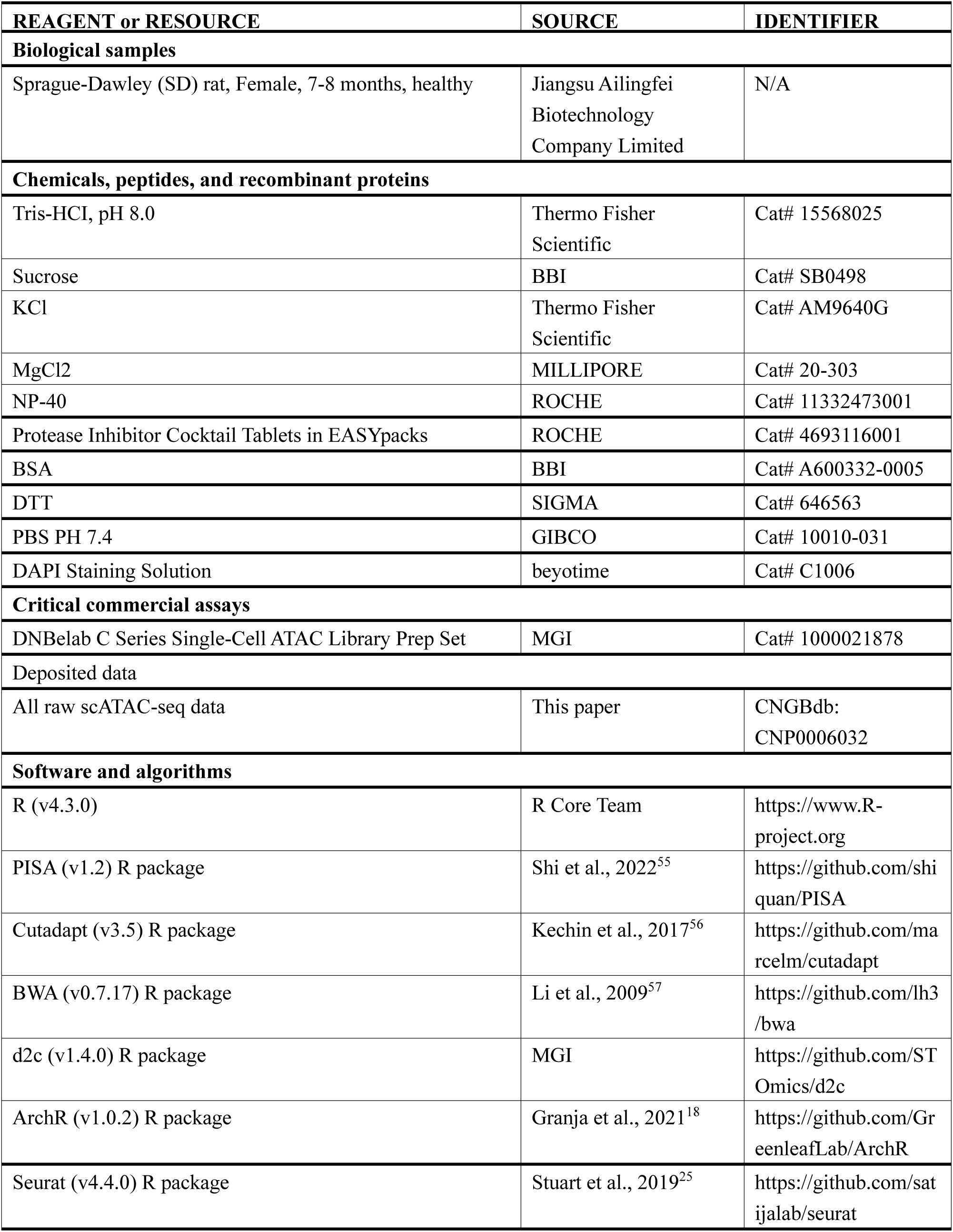
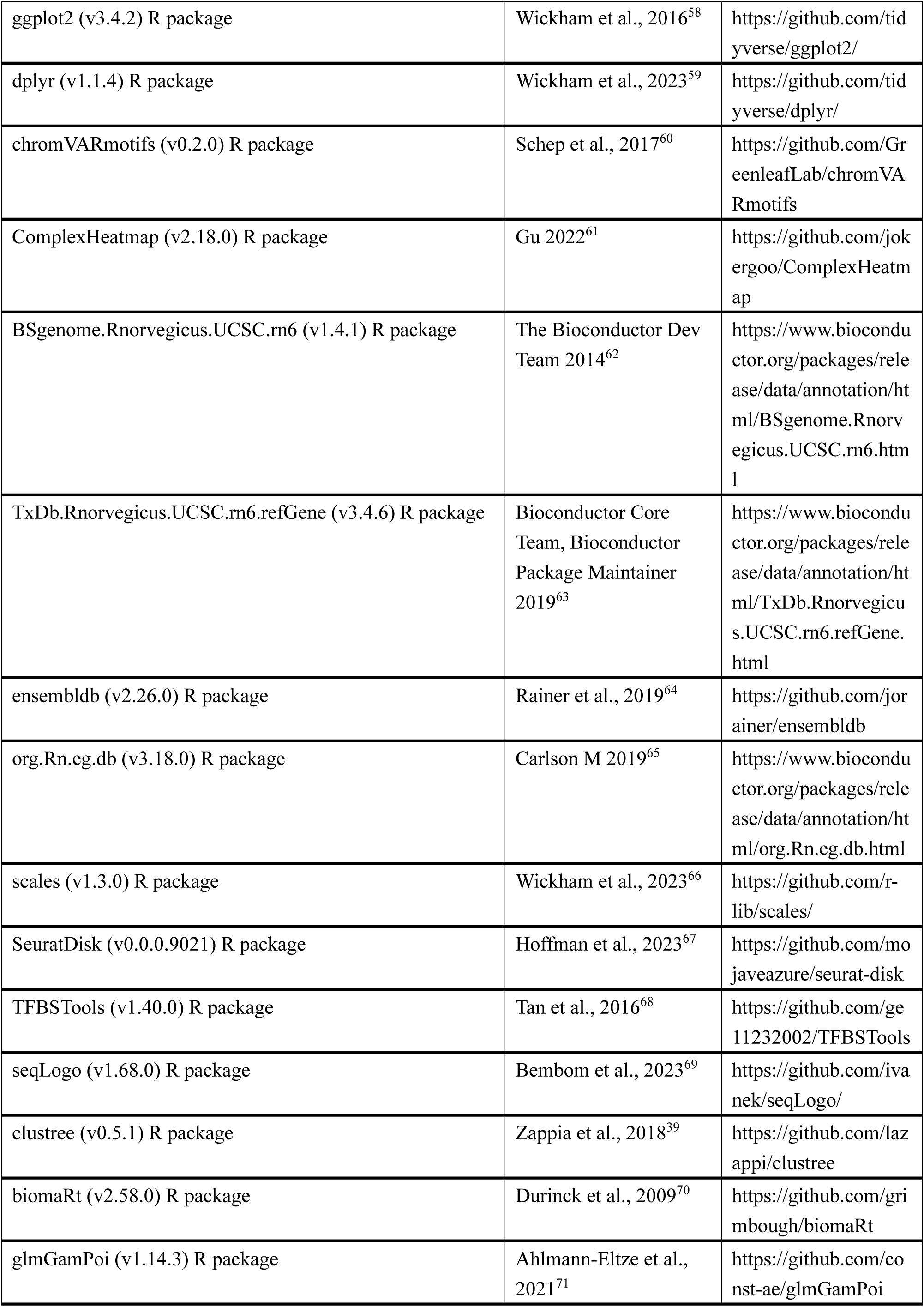
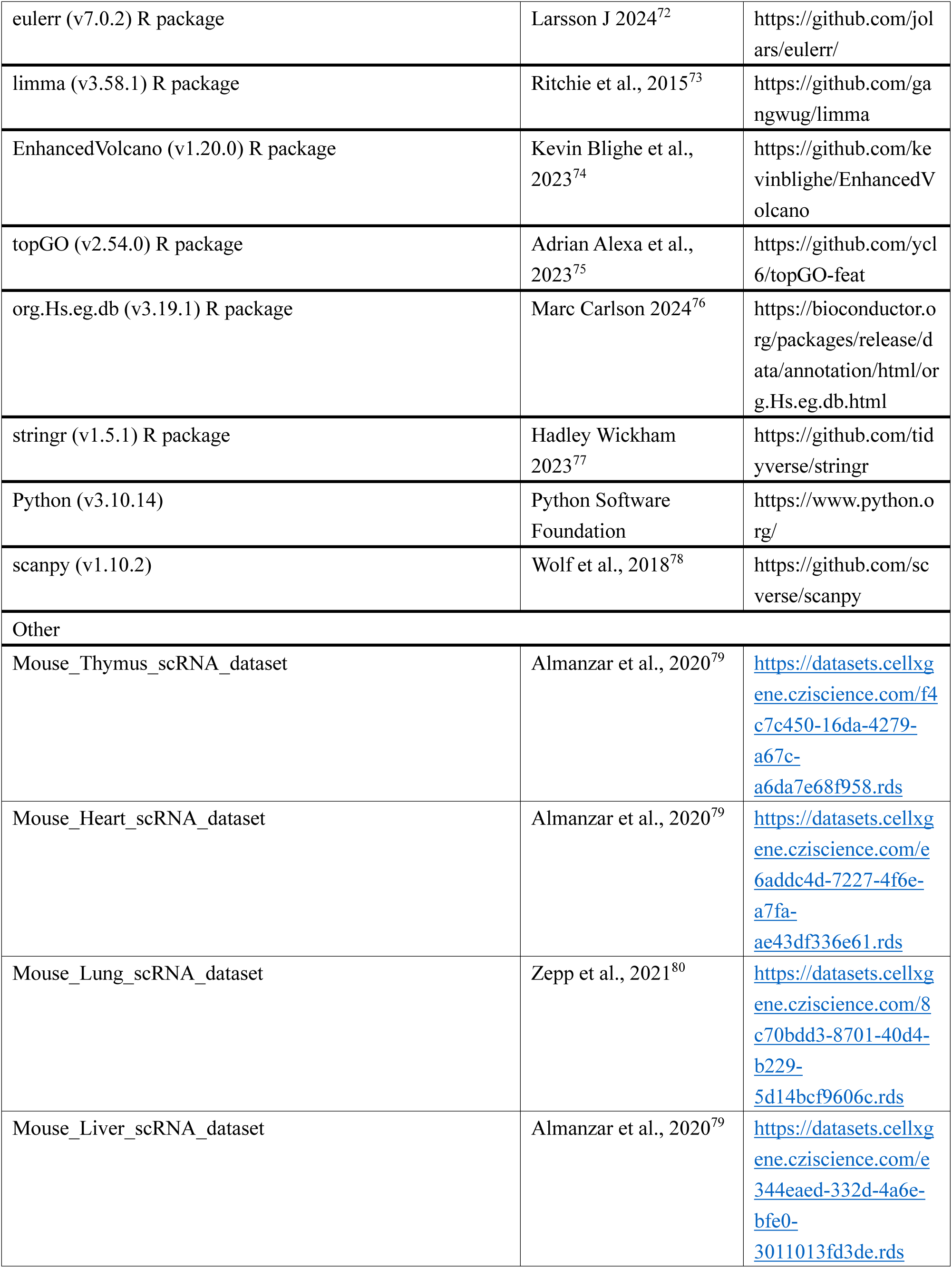
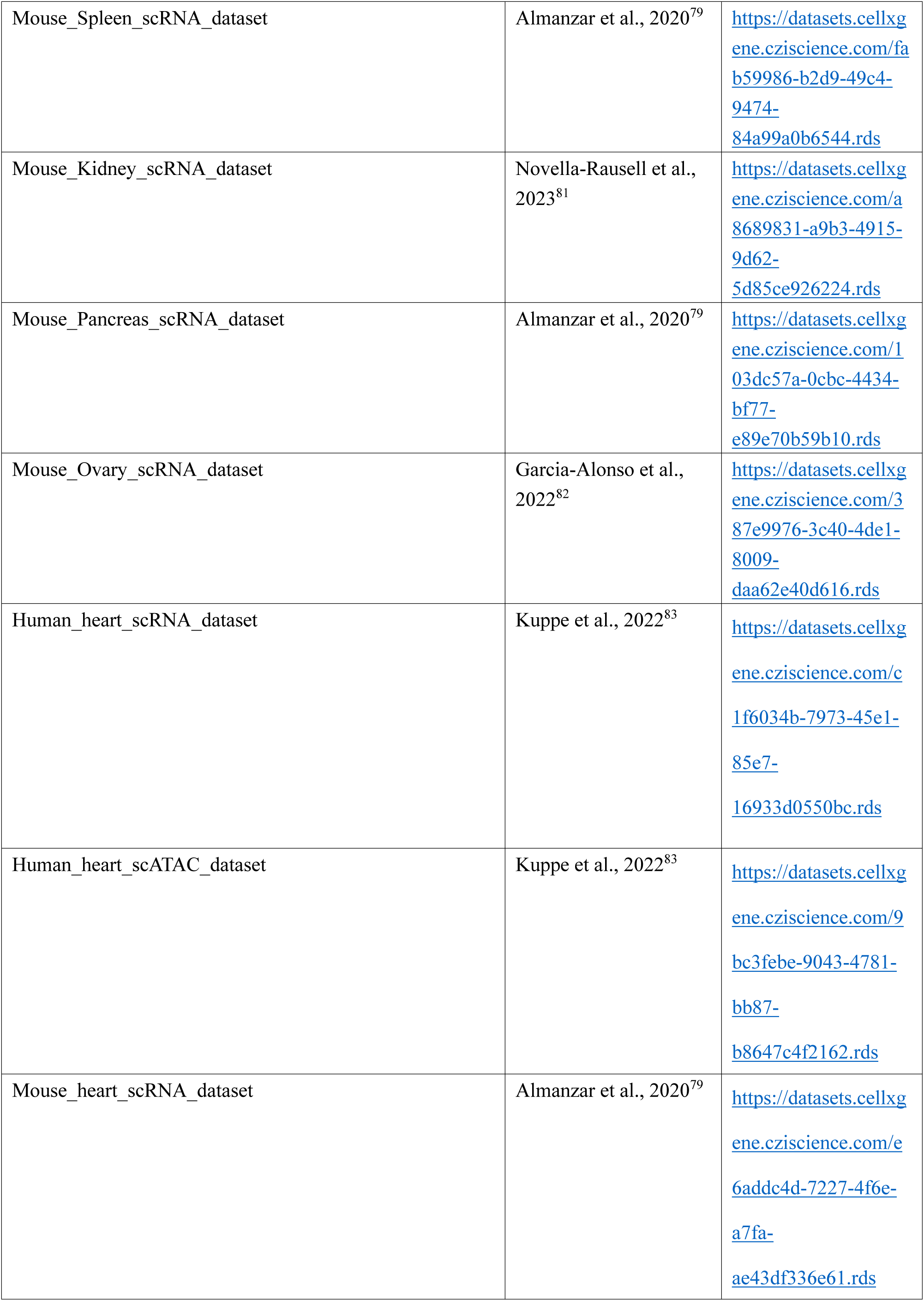
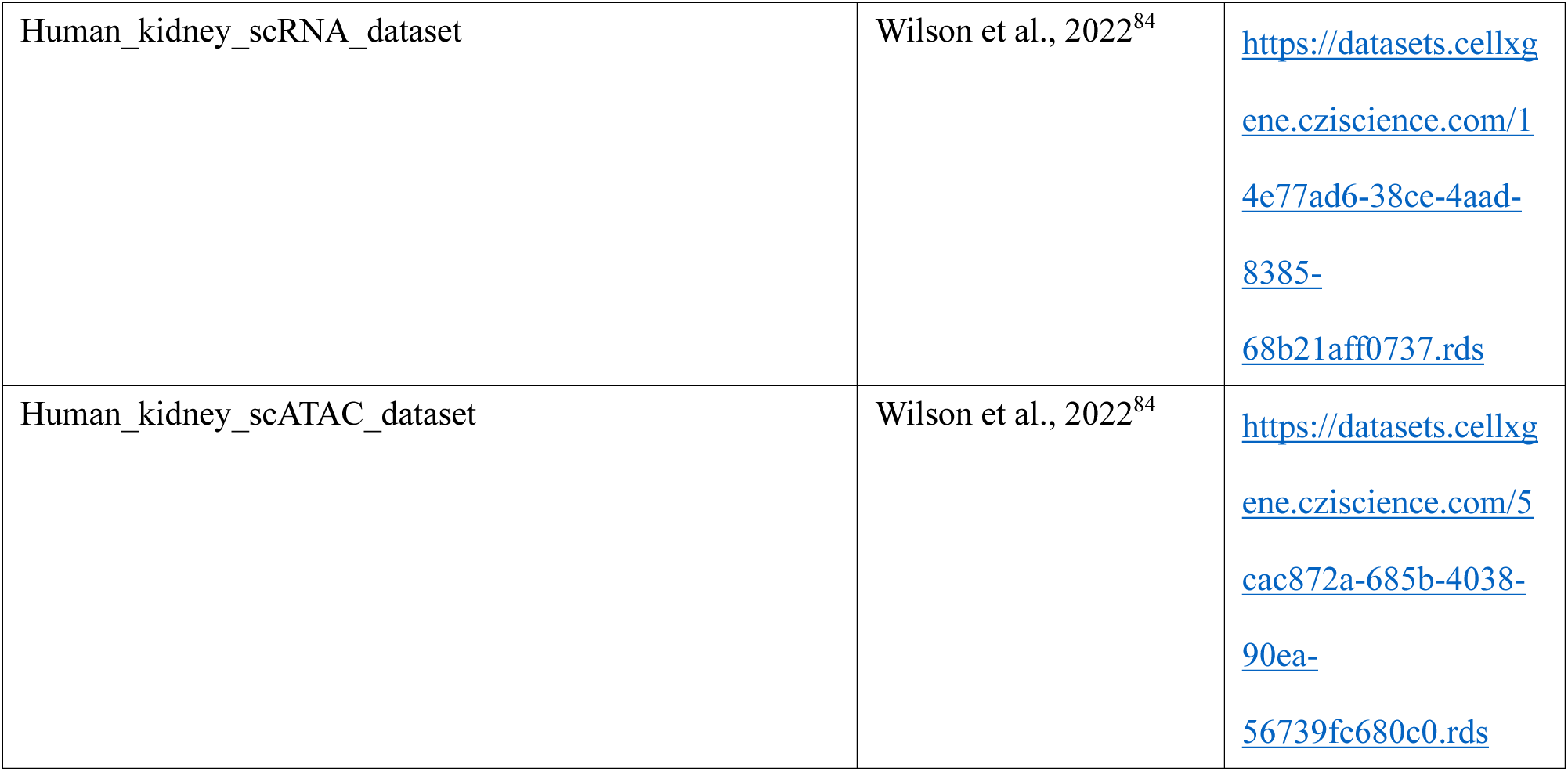
Key resources table

## Method details

### Tissue Dissection and Preservation

In this study, one healthy adult female Sprague-Dawley (SD) rat of 7-8 months of age was used. The animal was purchased from Jiangsu Ailingfei Biotechnology Company Limited. The dissection of rats was conducted by the Guangzhou Institute of Biomedicine and Health (GIBH), Chinese Academy of Sciences (CAS), in accordance with established protocols^85^. In summary, after executing the rats by carbon dioxide asphyxiation, we collected a total of nine organs, including thyroid, thymus, lung, heart, liver, spleen, pancreas, kidney, and ovary, and preserved them in liquid nitrogen tanks using cryopreservation tubes. The use of rats in the relevant experimental study was approved by the Institutional Review Board on the Ethics Committee of BGI (Permit No. BGI-IRB 23050-T1).

### Single nucleus ATAC sequencing

The single-cell experiment was divided into three steps.

Step one: Preparation of single-cell suspension. The preparation of single cell suspensions was conducted in accordance with the pre-established method^86^. Briefly, frozen tissue was cut into small pieces and transferred into a 2 mL KIMBLE Dounce Tissue Grinder (Sigma, #D8938-1SET) containing 2 mL of ice-cold homogenising buffer [20 mM Tris pH 8.0 (Thermo Fisher Scientific), 500 mM sucrose (BBI), 50 mM KCl (Thermo Fisher Scientific), 10 mM MgCl_2_ (MILLIPORE), 0.1% NP-40 (Roche), 1× protease inhibitor cocktail (Roche), and 1% nuclease-free BSA, and 0.1 mM DTT]. The tissues were homogenised by 15 strokes of the loose Dounce pestle, and the resulting homogenate was filtered through a 70 μM cell strainer (Falcon, # 352350). Subsequently, the filtered homogenate was subjected to 5 strokes of the tight pestle to facilitate the release of nuclei, which were then filtered once more through a 30 μM cell strainer (PLURISELECT, # 43-50030-03) and transferred to a 15 mL centrifuge tube. The filtered lysate was centrifuged at 500 g for 5 min at 4 °C. The pellet was then washed twice with 1 ml of ice-cold blocking buffer (1× PBS supplemented with 1% BSA), followed by another step of centrifugation at 500 g for 5 min at 4°C. Finally, the nuclei were resuspended in 50 μL of 1× PBS containing 1% BSA and counted with DAPI.

Step two: Construction of libraries. Single-nucleus ATAC-seq libraries were prepared using the DNBelab C Series Single-Cell ATAC Library Prep Set (MGI, #1000021878)^87^. Briefly, Chromatin-open regions were indexing in situ within the nucleus using Tn5 enzyme, after which the labelled nuclei are loaded into a DNBelab C4 microfluidic device for droplet encapsulation. This process is based on the principle of generating nanodroplets through the flow of two immiscible fluids (oil and water) within a microchannel. The droplets serve as discrete reaction chambers, each containing an individual cell nucleus and the requisite biochemical reagents^88^. Subsequently, the process entails PCR pre-amplification, emulsion breaking, bead collection, DNA amplification, and purification. In summary, we generated 25 single-cell ATAC libraries, with at least two technical replicates for each tissue sample.

Step three: Sequencing and alignment. All libraries were sequenced using the bipartite 50 sequencing protocol on the BGISEQ-500 and BGISEQ-T1 platform of the National Genebank of China (CNGB), with a minimum depth of 50,000 reads per nucleus for the libraries. Raw sequencing reads were demultiplexed using PISA^55^, adapters were removed using Cutadapt^56^, aligned to the rat genome (Rnor_6.0) using BWA^57^, and beads were called and merged using d2c(https://github.com/STOmics/d2c). The fragment file generated from each snATAC-seq library served as the basis for downstream analysis.

### Creating a custom archRGenome for Rat

In this study, the analysis of single-cell ATAC data was mainly conducted using the ArchR^18^ (v.1.0.2). The genome annotation was created using the *createGenomeAnnotation* function, specifying the rat genome as “BSgenome.Rnorvegicus.UCSC.rn6”. The gene annotation was generated with *createGeneAnnotation* function, using the TxDb (TxDb.Rnorvegicus.UCSC.rn6.refGene) and OrgDb (org.Rn.eg.db) objects to extract gene-related data, such as TSS, exons, and genes. Finally, the created genome and gene annotations were saved to an RData file. It should be noted that the custom ArchRGenome need to matche the reference genome used to generate the fragments file, which is crucial to avoid errors, such as issues in recognizing transcription start sites (TSS) when creating ArrowFiles.

### Preprocessing

The data preprocessing primarily involved three key steps:

First, the removal of low-quality nuclei. We used the *createArrowFiles* function to generate Arrow files from the fragment data. The data were filtered to exclude cells with fewer than 1000 unique nuclear fragments per cell or fewer than 4 TSS enrichment score per cell, as these metrics are crucial for ensuring that only nuclei with adequate chromatin accessibility and transcriptional activity are retained. During the quality control process, it was observed that many cells present in the thyroid samples exhibited a TSS value below 4, which typically indicates these cells are likely dead or dying, as their nucleosomes have begun to unravel. This unraveling can lead to random transposition events across the entire genome. Despite these cells having high levels of fragmentation (indicating potential chromatin activity), we were still removed from the analysis because they were classified as low-quality cells due to the low TSS enrichment score. This is notwithstanding the possibility that this could occur in certain biological states, such as dormant cells or specific cell types that naturally exhibit low levels of gene expression variability. Nevertheless, we were confident that we had taken the requisite precautions in sample processing. To guarantee the quality and accuracy of subsequent analyses, we retained the cells with high TSS enrichment.

Second, the elimination of potential doublets. we applied the *addDoubletScores* function to infer potential doublets, with the k parameter set to 10 to determine the number of nearest neighbors considered in the doublet detection process. Subsequently, we applied the *filterDoublets* function to remove doublets, with the filterRatio parameter set to 1. The filterRatio parameter controls the stringency of doublet removal; a higher filterRatio results in more cells potentially being identified and removed as doublets. For example, with a dataset of 5000 cells, the maximum number of cells that could be removed as doublets is computed as filterRatio * 5000^2 / 100000, which can be simplified to filterRatio * 5000 * 0.05.

Third, the exclusion of low-quality cell clusters. To enhances the accuracy of quality control, we used *subsetArchRProject* function to divide the entire dataset by organ for preprocessing. We used a for loop in R to perform same operations (like dimensionality reduction, clustering, visualization, marker gene analysis and heatmap generation) individually for each organ. For the dimensionality reduction and clustering, we employed *addIterativeLSI* and *addClusters* functions in ArchR, setting the parameters as follows: iterations at 3, resolution at c(0.2,0.4), varFeatures at 25,000, dimensions ranging from 1 to 30, and a resolution of 0.2 for clustering. For the visualization, we employed *addUMAP* in ArchR, setting the parameters as follows: nNeighbors at 60 and minDistat at 0.6. For the marker gene analysis, we employed *getMarkerFeatures* and *getMarkers* functions in ArchR, setting the parameters as follows: useMatrix at GeneScoreMatrix, groupBy at Clusters, testMethod at Wilcoxon and cutoff at FDR <= 0.01 & Log2FC >= 1. For the heatmap generation, we employed *plotMarkerHeatmap* functions in ArchR, setting the parameters as follows: cutoff at FDR <= 0.01 & Log2FC >= 1. During the viewing of the UMAP plot with Gene Scores Marker Heatmap, we manually identified low-quality clusters and removed them from the dataset based on the following rules:

a. The cluster did not express distinctly specific genes.
b. The number of cells of cluster less than 50.
c. The same cell cluster simultaneously expresses marker genes typical of multiple cell types and has a high doublet score.

After removal of low-quality cell clusters, we again performed the same operation as described above with the same parameters individually for each organ until the final clustering results in composite quality requirements.

### Annotation

To manually identify organ-specific cell types and states and to avoid confounding cell states due to batch effects caused by different organs, we employed *plotEmbedding* and *plotGroups* functions to visualize known marker genes of cell types individually for each organ, setting the parameters as follows: colorBy at GeneScoreMatrix, groupBy at Clusters. The list of known marker genes of cell types utilized in this study was derived from the aggregation of data from our previous investigation. Each cluster was meticulously annotated based on the established practices and insights provided by previous researchers^89^.

a. If a cluster expresses less than 3 markers related to a specific cell type with low expression, it is judged that the cluster does not belong to that cell type.
b. If multiple clusters co-express more than 3 markers related to the same cell type with high expression, it is judged that these clusters all belong to the same cell type.
c. If a cluster expresses multiple markers of different cell types and the first 10 marker genes of different cell types significantly mark the same cluster, the cluster is judged to be doublet and removed.

Additionally, we comprehensively considered the highly expressed gene profiles of each cluster while identifying each cell type. For the same cluster that unambiguously expresses marker genes of two cell types separately by contour region, the value of resolution in the *addClusters* function was appropriately increased to more accurately identify the cell type. We recommend trying multiple parameters and observing the cluster divisions when performing dimensionality reduction and cluster to balance the clustering granularity and biological interpretability.

### Label transfer

To help with cluster identity assignment, we used *addGeneIntegrationMatrix* function in ArchR to directly align cells from scATAC-seq with cells from scRNA-seq by comparing the scATAC-seq gene score matrix with the scRNA-seq gene expression matrix. This function converts the gene score matrix from the ArchR project into a Seurat object and uses *FindTransferAnchors* function from the Seurat^25^ package which allows you to perform CCA-based integration between the scATAC-seq data and the scRNA-seq data.

To compare predicted cell types from scRNA datasets with manually annotated cell types in scATAC datasets, we created a confusion matrix using the *confusionMatrix* function, calculated the similarity between the two sets of labels using the *jaccardIndex* function, added row and column annotations with the *HeatmapAnnotation* function, and then plotted the heatmap with customized aesthetics using the *heatmap* function.

Considering the current dearth of rat cross-organ single-cell RNA datasets, we employed data derived from mouse for integration in the present study. The download link for the mouse scRNA datasets used are as follows:

Thymus: https://datasets.cellxgene.cziscience.com/f4c7c450-16da-4279-a67c-a6da7e68f958.rds Heart: https://datasets.cellxgene.cziscience.com/e6addc4d-7227-4f6e-a7fa-ae43df336e61.rds

Lung: https://datasets.cellxgene.cziscience.com/8c70bdd3-8701-40d4-b229-5d14bcf9606c.rds Liver: https://datasets.cellxgene.cziscience.com/e344eaed-332d-4a6e-bfe0-3011013fd3de.rds

Spleen: https://datasets.cellxgene.cziscience.com/fab59986-b2d9-49c4-9474-84a99a0b6544.rds Kidney: https://datasets.cellxgene.cziscience.com/a8689831-a9b3-4915-9d62-5d85ce926224.rds

Pancreas: https://datasets.cellxgene.cziscience.com/103dc57a-0cbc-4434-bf77-e89e70b59b10.rds Ovary: https://datasets.cellxgene.cziscience.com/387e9976-3c40-4de1-8009-daa62e40d616.rds

All the above datasets were obtained from the online website CZ CELLxGENE: Discover^90,91^, with thanks to them for providing free access to convenient, standardized scRNA dataset downloads to facilitate the exploration and sharing of single cell datasets.

### Subcluster label assignment to full project

As described above, after identifying organ-specific cell types and low-quality clusters in each organ separately, we mapped these label matches back to the entire dataset for further analysis. For redoing dimensionality reduction and clustering, we used *addIterativeLSI* and *addClusters* functions, setting the parameters as follows: iterations at 2, resolution at 0.6, varFeatures at 25,000, dimensions ranging from 1 to 30, and a resolution of 0.2 for clustering.

To visualize the expression of marker genes for each cell type in each organ, we employed the *dotplot* function in Scanpy^78^. We first need to convert the GeneScoreMatrix from ArchR project into a Seurat object. We got the gene score matrix from ArchR project using the *getMatrixFromProject* function and created Seurat object using the *CreateAssayObject* and *CreateSeuratObject* function. The Seurat object was normalized (LogNormalize method) and variable features were identified (vst method with 2000 features) using *NormalizeData* and *FindVariableFeatures* functions. It was then saved in h5Seurat format using *SaveH5Seurat* function and converted into an h5ad format using *Convert* function, which is compatible with AnnData, often used in Python for further single-cell RNA-seq analysis. Additionally, the metadata from the ArchR project was extracted using *getCellColData* function and saved as a CSV file. In the Scanpy analysis, we used the default parameters to identify differentially expressed genes between cell subtypes and then filtered these results by a minimum fold change of 1. We subset each organ data to generated the expression of marker genes using *sc.pl.dotplot* function.

To compute hierarchical clustering of cell subtypes, we employed *sc.tl.dendrogram* functions with using “complete” linkage and optimal ordering to understand the hierarchical relationships between different cell subtypes.

### Peaks calling

We created pseudo-bulk replicates, a bulk ATAC-seq experiment, allowing for more robust downstream analyses by reducing noise and enabling statistical comparisons, based on major cell types in the dataset using *addGroupCoverages* function in ArchR. To call peaks using MACS2^92^, we utilized *addReproduciblePeakSet* function to generate reproducible peak set across cells grouped by major cell types in ArchR.

### Motif Enrichments

To determine which transcription factors (proteins that bind to specific DNA sequences to regulate gene expression) are responsible for binding events, we utilized *addMotifAnnotations* function to add motif information to the ArchR project, setting the parameters as follows: motifSet at cisbp^93^, species at mus musculus. Although rat-specific motifs are ideal, the limited availability justifies the use of mouse motifs. Many TF binding sites are conserved across closely related species, so using mouse motifs can still provide meaningful insights.

For the motif enrichments analysis, we employed *getMarkerFeatures*, *peakAnnoEnrichment* and *plotEnrichHeatmap* functions in ArchR, setting the parameters as follows: useMatrix at PeakMatrix, groupBy at cell subtypes or organs, testMethod at wilcoxon and cutoff at FDR <= 0.1 & Log2FC >= 0.5.

For identifying and visualizing the most enriched TFs associated with chromatin accessibility in different major cell types, we identified marker genes (via GeneScoreMatrix) and motif enrichments (via PeakMatrix) across major cell types and merge gene scores with motif enrichment data, linking TF activity to specific gene expression patterns.

### Motif Footprinting

We began by extracting motif positions within the genome using *getPositions* function. These positions are where specific TF motifs are predicted to occur. To enhance the signal for foot printing analysis, we calculated group coverages for different organs using *addGroupCoverages* function. To visualize the accessibility profile around the motif sites, we used *getFootprints* and *plotFootprints* functions with bias subtraction. To visualize sequence logos for specific motifs, we employed TFBSTools^94^ Package to converts position weight matrices (PWMs) into probability matrices suitable for visualization with seqLogo (<u>10.18129/B9.bioc.seqLogo</u>).

### Analysis imumune cells across organs

To identify and analysis of immune cell diversity and regulatory elements across organs, we first extracted immune cells from the full dataset and recalled specific peaks for cell subtypes (consistent as described in the previous methods but grouped by cell subtype). Dimensionality reduction was then performed using *addIterativeLSI* function, setting the parameters as follows: iterations at 3, resolution at c(0.4,0.6), varFeatures at 25,000, dimensions ranging from 1 to 25. To correct for batch effects between different organs, we used *addHarmony* function in ArchR. Next, clustree package was used to optimise and visualise the clustering process, and finally the analysis was further visualised in low-dimensional space using *addUMAP* function with nNeighbors at 70 and minDist at 1.4.

We quantify the accessibility (gene score activities) of genes known to be markers of different stages in T cell development to selecting a predefined sequence of cell clusters. By using the *addTrajectory* function in ArchR, we integrated these clusters into a developmental trajectory named “T_cell_development.” This trajectory is visualized using UMAP, with cells colored based on their position in the trajectory. To visualize of how specific genes are regulated as T cells progress through developmental stages, we used *getTrajectory* and *plotTrajectoryHeatmap* functions, setting the parameters as follows: useMatrix at GeneScoreMatrix, smoothWindow at 5, labeltop at 20. To analysis of T cell development by integrating motif activity (using ChromVAR) with the gene score trajectory data, we first calculated the deviation matrix for motif activities using *addBgdPeaks* and *addDeviationsMatrix* function, and then extracted motif trajectory using *getTrajectory* function with MotifMatrix. Next, we identified motifs that match gene activity patterns by correlating the gene score trajectory with the motif activity trajectory using *correlateTrajectories* function, and finally heatmaps were generated to visualize the combined trajectories using *plotTrajectoryHeatmap* function.

To explore the regulatory landscape of T cell differentiation, we began by refining cluster labels to distinguish CD4^+^ and CD8^+^ T cells within cluster 20. We then defined genomic regions and loops around T cell markers with *getCoAccessibility* function. Finally, we visualized these elements using *plotBrowserTrack* function, integrating feature annotations, and co-accessibility data.

### Cross species integration

The process of cross-species data integration can be divided into three steps.

Step one: Data preprocessing. For the rat dataset, we extracted the GeneScoreMatrix from an ArchR project using *getMatrixFromProject* function and converted it into a Seurat object using *CreateAssayObject* and *CreateSeuratObject* functions. To ensure consistency of gene names across species datasets, we converted gene symbols in the Seurat object to Ensembl IDs using the *bitr* function from the clusterProfiler^95^ package, leveraging the org.Rn.eg.db database. To ensure comparability of cell types in cross-species data integration and comparative analyses, we screened for homologous cell types across species and down-sampled according to cell type to ensure that the number of cells of each cell type in the analyses is in a reasonable range (e.g., a minimum of 50 and a maximum of 1000).

Step two: Homologous substitution of gene names. We connected to the Ensembl database using the biomaRt^70^ package and converted rat and mouse gene symbols to their human homologs. Step three: Cross species integration. We began by compiling the data into a list of Seurat objects. To normalize and standardize the data, we applied the *SCTransform* function with the glmGamPoi method to each dataset in the list, ensuring that all variable genes were retained (return.only.var.genes = F). Next, we selected 3,000 integration features across the datasets using the *SelectIntegrationFeatures* function, which identifies the most consistent and variable genes for integration. We then prepared the data for integration using *PrepSCTIntegration* and identified anchors across the datasets with *FindIntegrationAnchors*, using the selected features and the first 30 principal components (dims = 1:30) to align the datasets. We integrated the data using the *IntegrateData* function, which combined the datasets into a single Seurat object normalized with the SCT method. After integration, we reduced the dimensionality of the data using PCA with *RunPCA*, and then visualized it in a lower-dimensional space using UMAP with *RunUMAP*. Finally, we identified cell clusters by *FindNeighbors* and clustering them with the Louvain algorithm (*FindClusters*), setting the resolution to 0.3 to control the cluster size. To identify the shared and unique DEGs among human, mouse, and rat, we subsetted into separate datasets for Human, Mouse, and Rat and down sampled each dataset to 200 cells per cell type to ensure comparable cell numbers across species. we then identified DEGs for each cell type within each species using the *FindAllMarkers* function with default parameters. To visualize represent the overlap of DEGs among Human, Mouse, and Rat, we generated Venn diagram using the eulerr package (https://github.com/jolars/eulerr). To perform differential expression analysis across species (Human vs. Mouse, Human vs. Rat, Mouse vs. Rat), we ued the *FindMarkers* function with default parameter. Volcano plots was generated using the EnhancedVolcano package (https://github.com/kevinblighe/EnhancedVolcano). Gene Ontology (GO) enrichment analysis was performed using topGO package (https://bioconductor.org/packages/release/bioc/vignettes/topGO/inst/doc/topGO.pdf).

The download link for the human or mouse scRNA/scATAC datasets used for Cross species integration are as follows:

Human_heart_scRNA-seq:

https://datasets.cellxgene.cziscience.com/c1f6034b-7973-45e1-85e7-16933d0550bc.rds Human_heart_scATAC-seq:

https://datasets.cellxgene.cziscience.com/9bc3febe-9043-4781-bb87-b8647c4f2162.rds Mouse_heart_scRNA-seq:

https://datasets.cellxgene.cziscience.com/e6addc4d-7227-4f6e-a7fa-ae43df336e61.rds Human_kidney_scRNA-seq:

https://datasets.cellxgene.cziscience.com/14e77ad6-38ce-4aad-8385-68b21aff0737.rds Human_kidney_scATAC-seq:

https://datasets.cellxgene.cziscience.com/5cac872a-685b-4038-90ea-56739fc680c0.rds

All the above datasets were obtained from the online website CZ CELLxGENE: Discover, with thanks to them for providing free access to convenient, standardized scRNA dataset downloads to facilitate the exploration and sharing of single cell datasets.

## Data availability

The data supporting the findings of this study have been deposited into CNGB Sequence Archive (CNSA)^96^ of China National GeneBank DataBase (CNGBdb)^97^ with accession number CNP0006032.

## Code availability

We have described the process of analyzing the data for this study in detail in the methods section. If you have questions about data analysis or are looking for the full code, please contact by mail at.

The software repositories and documentations are accessible through the links provided below: ArchR (https://www.archrproject.com/bookdown/index.html), Seurat (https://satijalab.org/seurat/), Scanpy (https://scanpy.readthedocs.io/en/stable/tutorials/index.html). Part of the code is referenced in the following link:https://github.com/winstonbecker/scCRC_continuum?tab=readme-ov-file, https://github.com/GreenleafLab/scScalpChromatin/blob/main. Thanks to them for sharing such great code to boost the community.

## Acknowledgements

We thank all members of our teams and the China National GeneBank (CNGB) for their support, and we are especially grateful to Dr. Duoyuan Chen, Dr. Xi Dai and Dr. Shijie Hao of BGI for their helpful comments in cross-species analysis. This work was supported by the Shenzhen Key Laboratory of Single-Cell Omics (ZDSYS20190902093613831).

## Author contributions

C.L. and Y.Y. designed the project and experiments. Y.Y., S.D. and Q.D. conducted snATAC-seq experiments. W.M. processed the raw sequencing data. R.L. performed the data analysis and wrote the manuscript. C.L. participated in the supervision of this research. R.L., Y.Y., C.L., L.L. and P.G. revised the manuscript. All authors have read and approved the final manuscript. The remaining authors declare no competing interests.

## Supplemental information

**Document S1. Figures S1–S5**

**Table S1. The main marker genes were used for annotation in this paper, related to Figure S1 and S2**

**Data S1. The metadata of scATAC dataset in this paper, related to Figure 1 and S1**

**Data S2. The data frame that contains the UMAP coordinates for each cell in the ArchR project, related to Figure 1**

**Data S3. The data frame contains information about top20 genes identified for each cell subtype, related to Figure 2**

**Data S4. The data frame contains the results of specific TF binding motifs are significantly enriched in the accessible chromatin regions (peaks) associated with different cell subtypes, related to Figure 3**

**Data S5. The data frame contains the results of specific TF binding motifs are significantly enriched in the accessible chromatin regions (peaks) associated with different major cell types, related to Figure 3**

**Data S6. The data frame contains the results of specific TF binding motifs are significantly enriched in the accessible chromatin regions (peaks) associated with different organs, related to Figure 3**

**Data S7. A comprehensive dataset that consolidates differential expression analysis results across multiple subclasses and species., related to Figure 5**

