## Supplementary material for "Single-nucleus multiple-organ chromatin accessibility mapping in the rat": Document S1.docx

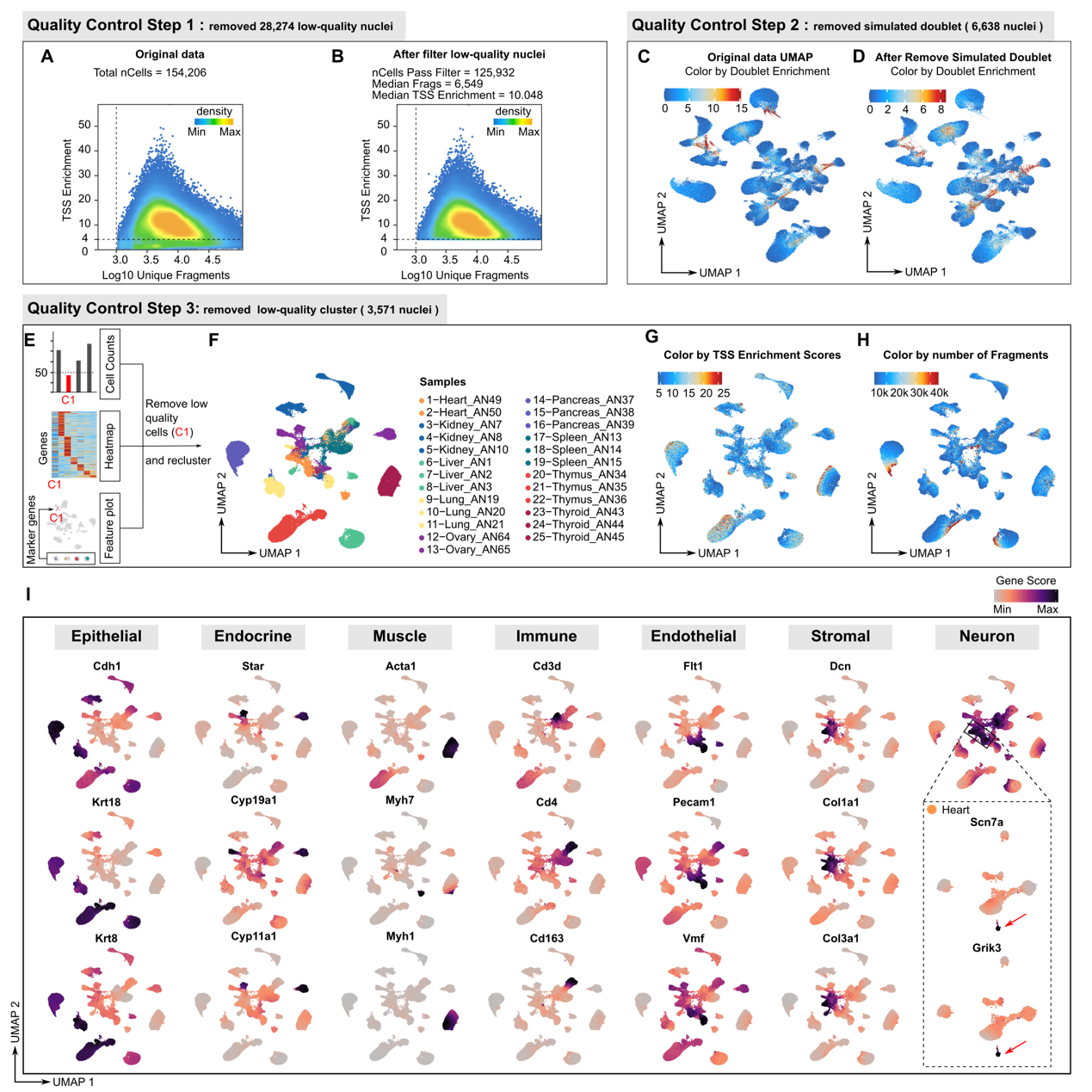

**Figure S1.** Quality control and analysis workflow (related to Figure 1). **A**. Density plot shows the relationship between the log10 unique nuclear fragment count and transcription start site (TSS) enrichment before filtering. **B.** Density plot shows the relationship between the log10 unique nuclear fragment count and transcription start site (TSS) enrichment after filtering. The plot reveals the distribution of nuclei that were filtered and met the quality thresholds, of which there were 125,932 in total. **C**. UMAP plot shows all nuclei before removal of simulated doublets, colored by doublet enrichment indicating simulated multiplets, with high scores (red) indicating low-quality single nuclei. **D**. Same as C, but for all nuclei after removal of simulated doublets. **E**. Schematic illustration of the process of identifying low-quality clusters. **F.** UMAP plot shows all nuclei after removing low-quality clusters, colored by sample. **G**. Same as F, but colored by TSS enrichment scores. **H**. Same as F, but colored by number of fragments. **I**. UMAP plot illustrates the gene activity score of known marker genes across major cell types. Each row corresponds to a specific gene, while each column represents a different cell type category. Particularly, with an inset focusing on neuron clusters and an arrow indicating the specific expression pattern of *Scn7a* and *Grik3* in relation to heart tissue.

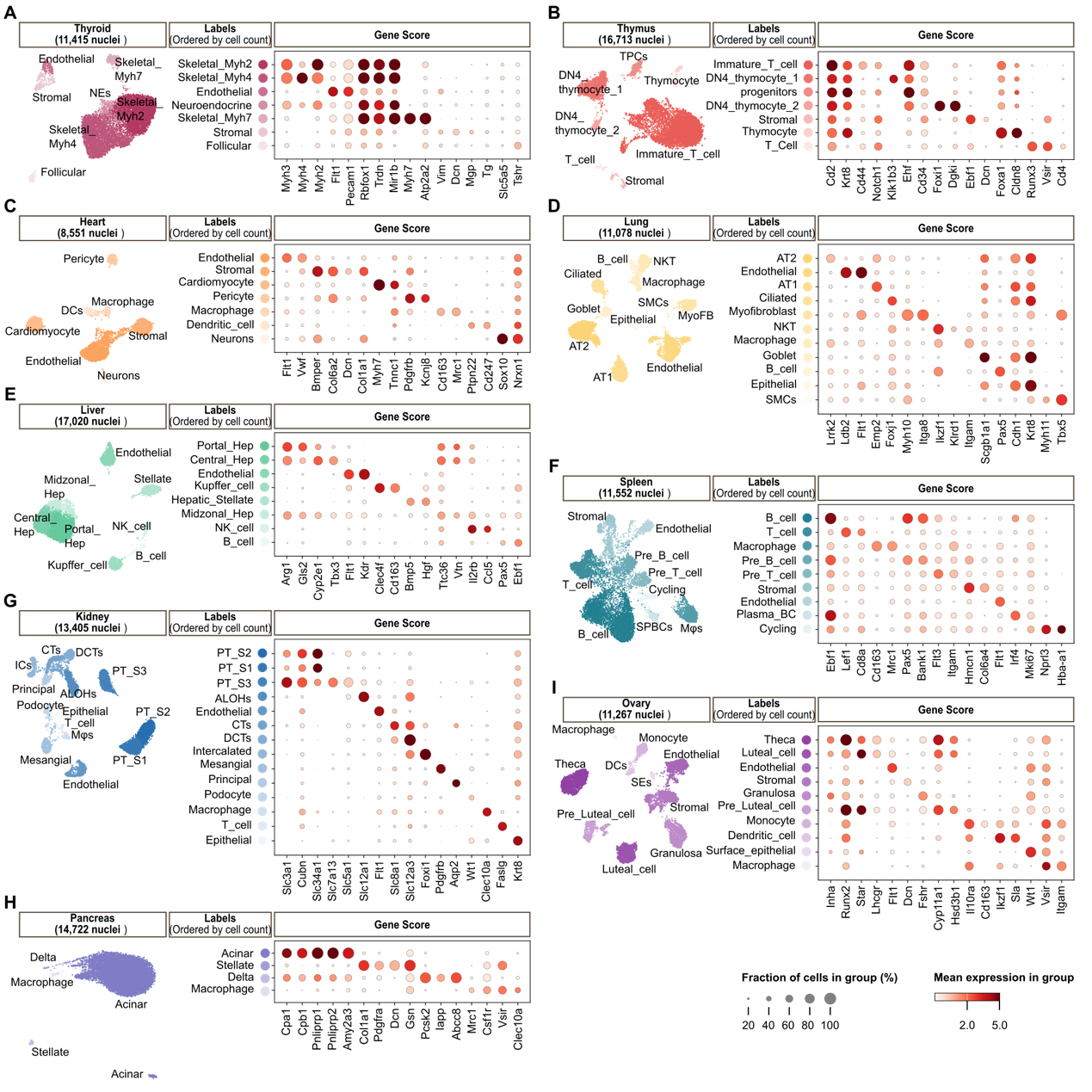

**Figure S2.** A comprehensive overview of cell type-specific gene expression patterns for each organ (related to Figure 2). **A-I.** Each panel shows UMAP plots (left) and corresponding dot plots (right) for each organ. The UMAP plot visualizes clusters of different cell types, the labeling is based on the number of cells of the cell type using over-coloring, the darker the color, the greater the number of cells. The dot plot displays the expression levels of marker genes across cell types, the size of the dot corresponds to the percentage of cells expressing the genes in each cell type, the color represents the average expression level.

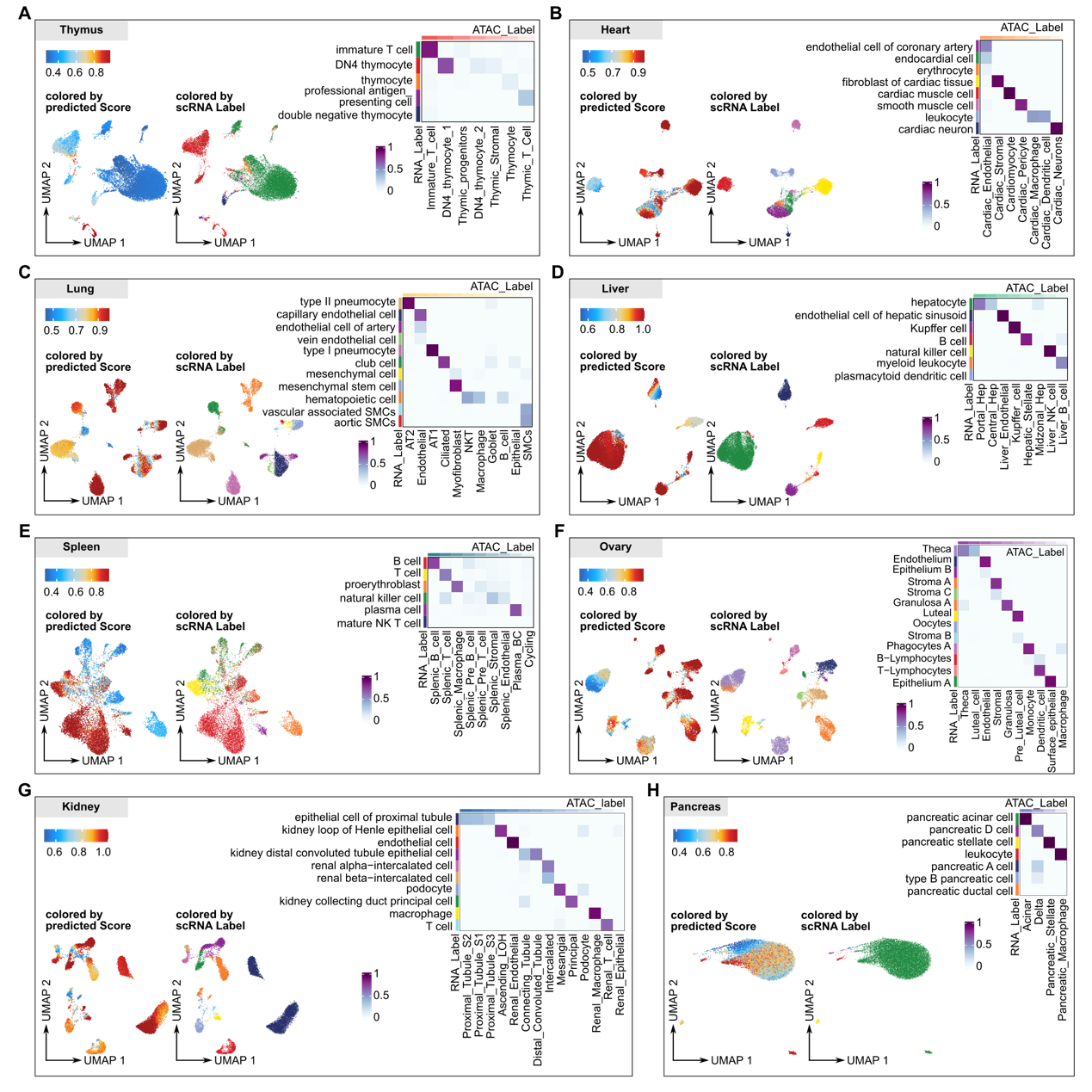

**Figure S3.** Integration of single-cell RNA sequencing and single-nucleus ATAC sequencing data for each organ (related to Figure 2). **A-H.** Each panel represents a different organ, displaying two UMAP plots alongside a heatmap. The left UMAP is colored by the predicted score of cell types from scRNA-seq data. The predicted scores represent the accuracy of the group assignment in the RNA cells. Lower scores represent ambiguous predictions, and higher scores represent accurate predictions. The right UMAP is colored by the predicted labels of cell types from scRNA-seq in the snATAC dataset. The heatmap compares the cell types from the snATAC-seq data with the cell type labels from the scRNA-seq data using a Jaccard index score. The cell types of the snATAC-seq are manually identified by gene activity scores, which aggregate the chromatin accessibility around a gene body. Higher values (darker colors) on the diagonal indicate greater overlap between RNA and ATAC-seq derived labels, demonstrating accurate cell type annotation.

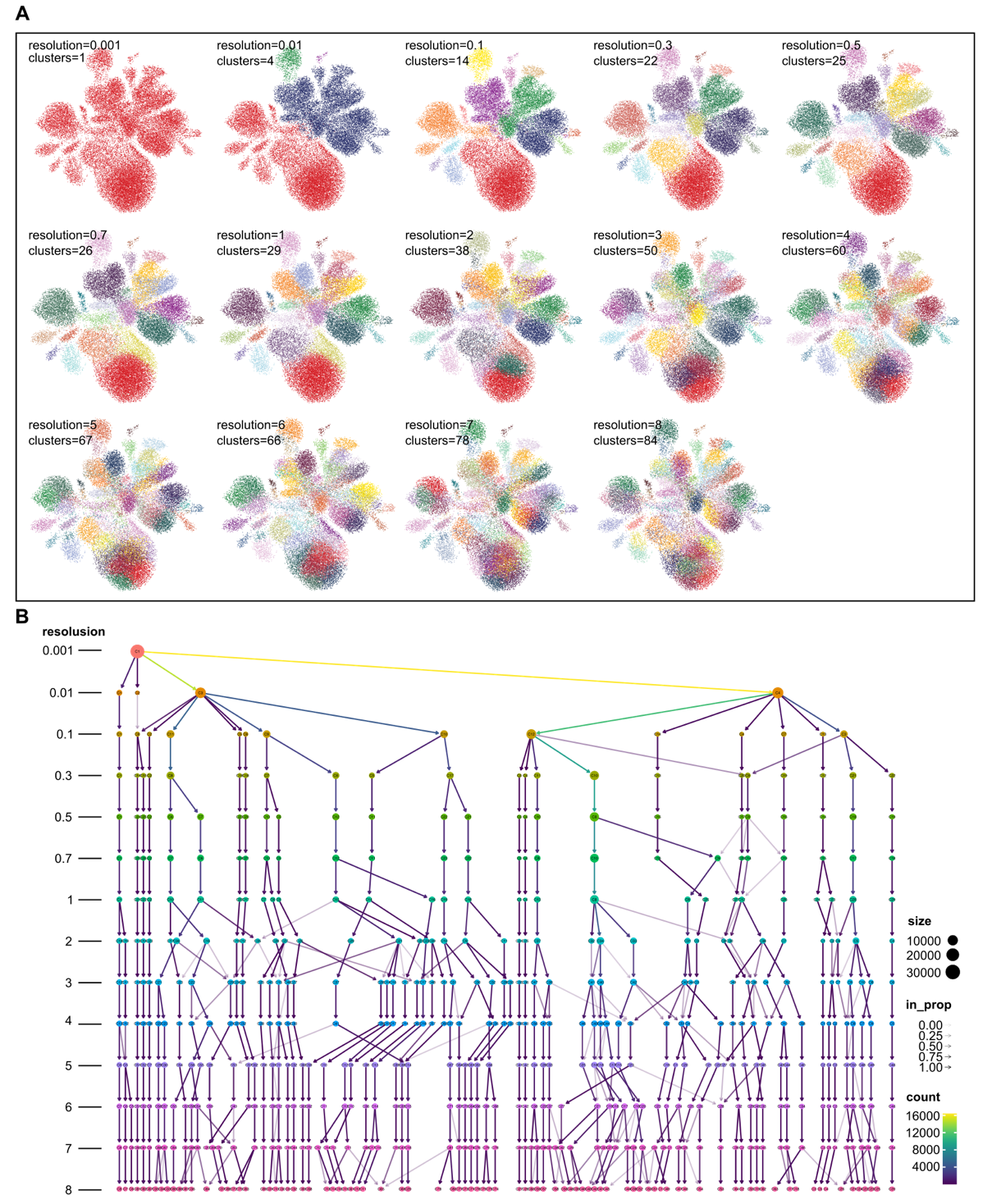

**Figure S4.** The balance between granularity and biological interpretability (related to Figure 4)**.** **A.** UMAP plots of clusters at different resolutions, each generated using a different resolution parameter for clustering. **B.** Clustering tree across resolutions. This illustrates the relationships between clusters at different resolutions. Each node in the tree represents a cluster, and the lines connecting nodes represent how clusters at lower resolutions break down into finer clusters at higher resolutions. The nodes are colored according to the value of, and their size is determined according to the number of cells they represent. Edges are colored according to the number of cells (from blue for few to yellow for many). Transparency is adjusted according to the ratio, with thicker lines indicating edges that are more important to the higher resolution cluster. Cluster labels are randomly assigned by the algorithm.

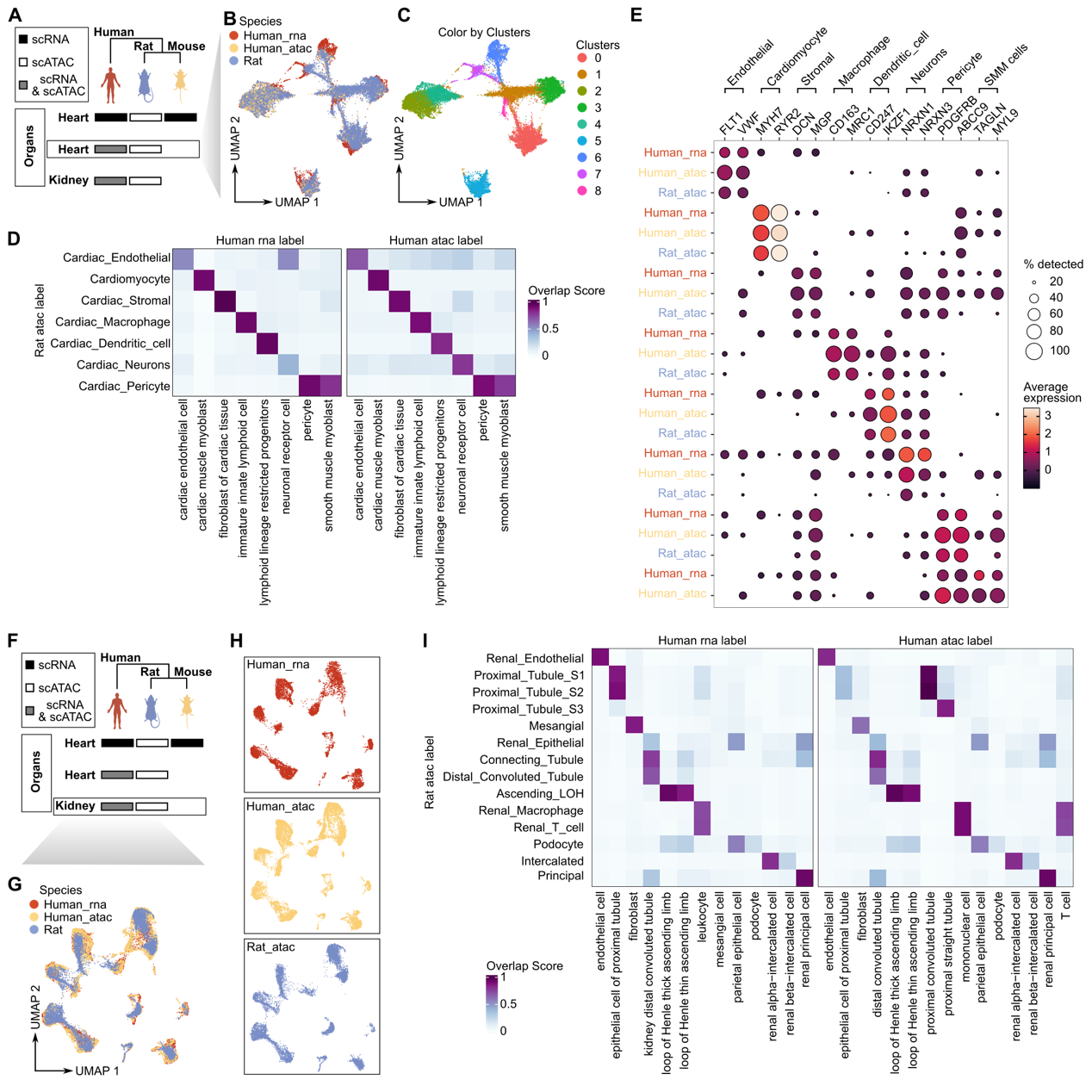

**Figure S5.** Cross-species integration of multi-omics datasets in heart and kidney (related to Figure 5). **A**. Overview of data types (scRNA-seq and scATAC-seq) collected from heart and kidney tissues in human, rat, and mouse. **B.** UMAP plot shows species-specific clustering. **C.** UMAP plot shows the clustering of single cells from the integrated dataset, colored by distinct clusters. **D**. Heatmaps shows the overlap score between rat and human on cell types for both scRNA-seq and scATAC-seq data in the heart. **E**. Dot plot displays the gene expression levels of known marker genes of heart tissue in different clusters, split by species. The size of the dot corresponds to the percentage of cells expressing the genes in each cluster, the color represents the average expression level. **F.** Same as A, but focus on kidney. **G.** UMAP plot shows species-specific clustering. **H.** UMAP plot shows species-specific clustering, split by species. **I**. Heatmaps shows the overlap score between rat and human on cell types for both scRNA-seq and scATAC-seq data in the kidney.
